# Subcortical Segmentation of the Fetal Brain in 3D Ultrasound using Deep Learning

**DOI:** 10.1101/2021.09.29.462430

**Authors:** Linde S. Hesse, Moska Aliasi, Felipe Moser, the INTERGROWTH-21^st^ Consortium, Monique C. Haak, Weidi Xie, Mark Jenkinson, Ana I.L. Namburete

**Affiliations:** Institute of Biomedical Engineering, University of Oxford, United Kingdom; Department of Obstetrics and Fetal Medicine, Leiden University Medical Center, The Netherlands; Visual Geometry Group, Department of Engineering Science, University of Oxford, United Kingdom; Wellcome center for Integrative NeuroImaging, FMRIB, University of Oxford, United Kingdom; Australian Institute for Machine Learning (AIML), Australia; South Australian Health and Medical Research Institute (SAHMRI), Australia; Department of Computer Science, University of Oxford, United Kingdom

**Author notes:** Corresponding author *Email address:* (Linde S. Hesse).

**Keywords:** Ultrasound, Segmentation, Fetal, Brain, Subcortical, Convolutional Neural Network, Few-Shot Learning, Deep Learning

## Abstract

The quantification of subcortical volume development from 3D fetal ultrasound can provide important diagnostic information during pregnancy monitoring. However, manual segmentation of subcortical structures in ultrasound volumes is time-consuming and challenging due to low soft tissue contrast, speckle and shadowing artifacts. For this reason, we developed a convolutional neural network (CNN) for the automated segmentation of the choroid plexus (CP), lateral posterior ventricle horns (LPVH), cavum septum pellucidum et vergae (CSPV), and cerebellum (CB) from 3D ultrasound. As ground-truth labels are scarce and expensive to obtain, we applied few-shot learning, in which only a small number of manual annotations (n = 9) are used to train a CNN. We compared training a CNN with only a few individually annotated volumes versus many weakly labelled volumes obtained from atlas-based segmentations. This showed that segmentation performance close to intra-observer variability can be obtained with only a handful of manual annotations. Finally, the trained models were applied to a large number (n = 278) of ultrasound image volumes of a diverse, healthy population, obtaining novel US-specific growth curves of the respective structures during the second trimester of gestation.

## 1. Introduction

During pregnancy, several subcortical structures in the fetal brain are assessed with ultrasound (US) imaging. Especially earlier in gestation, when the fetal skull is not yet fully calcified, the US beam can penetrate the skull and visualize the subcortical structures. The abnormal development of subcortical structures can be a potential sign of a severe neurological condition and, as such, it is important to monitor their development during gestation. For example, an enlarged ventricular cavity can be a sign of a central nervous system (CNS) abnormality [1] whereas the inability to visualize the CSP after 17-20 gestational weeks (GWs) may be indicative of underdevelopment of the corpus callosum, which is a fiber track connecting the two cerebral hemispheres [1–3]. Brain development can be studied in detail with targeted fetal neurosonography [4], however, this is only performed in fetuses at high-risk for CNS abnormalities and is not part of routine obstetric examinations. For this reason, efforts should be made to develop analysis methods that can improve subcortical assessment during routine pregnancy monitoring.

Subcortical structure development is best quantitatively assessed using 3D volumetric information. However, to date, most studies analyzing these structures during gestation only use in-plane measurements obtained from 2D US [5, 6]. These measurements are routinely acquired during a basic examination of the fetal brain [1] but only provide a limited representation of the anatomical development and morphology. For this reason, it is desirable to use 3D US to obtain volumetric measures of subcortical development. As manually annotating several structures is not feasible in routine clinical practice, automated subcortical segmentation methods for 3D US could facilitate this analysis and may provide new insights into in utero subcortical development.

Structural segmentation in fetal brain US is a challenging task due to low soft tissue contrast, reverberation artifacts and the characteristic presence of speckle. Consequently, precise structural boundaries can be hard to distinguish, resulting in high inter- and intra-observer variability in manual annotations. Since manual segmentation is not a task usually performed in clinical practice, even trained ultrasonographers can have difficulty in accurately segmenting subcortical structures in 3D US volumes. An additional challenge of US volumes acquired with a free-hand scanning protocol, as is typical at the bedside, is the varying position of the fetal brain due to the unpredictable fetal position in the womb as well as movement of the transducer relative to the fetal head. Furthermore, due to interactions of the US beam with the fetal skull, typically only the cerebral hemisphere distal to the US transducer is well visible in the US volumes.

Recently, it has been shown that deep learning methods can be successfully applied to other segmentation tasks in 3D US volumes of the fetal brain [7–9], resulting in higher performance than traditional image analysis methods. However, due to the difficulty of obtaining manual annotations for subcortical structures, a key barrier to applying deep learning methods to this task is obtaining sufficient ground-truth annotations for training. Therefore, to the best of our knowledge, only two previous studies have applied deep learning to subcortical structure segmentation in 3D US, which will be discussed in more detail in Section 2.1.

To overcome the need for a large manually annotated dataset, few-shot learning can be used, in which only a small number of manual annotations are used to train a convolutional neural network (CNN). Several fewshot learning approaches have been proposed for segmentation tasks in the medical image domain [10–12], such as training a CNN with pseudo-labels obtained from few annotated 2D slices [12] or with additionally generated labels by a generative adversarial network [11]. These studies thus show that good segmentation performance can be obtained using only a very limited amount of voxel-wise manual annotation.

In this work, we will use few-shot learning to develop a deep-learning based method for the segmentation of several subcortical structures in 3D US. To the best of our knowledge, few-shot learning has not been applied for this task, and, as such, this will be the first study exploring subcortical segmentation of the fetal brain in a low-data regime. Specifically, we will compare two types of ground-truth labels for training, where both have been obtained using only a few manual annotations: (1) a small number of individually annotated US volumes (referred to as *expert labels*), and (2) a large number of weakly labelled US volumes obtained by propagating annotated template images (referred to as *atlas labels*).

Additionally, we investigate the impact of image alignment to a common coordinate system as a pre-processing step. As the anatomical orientation of the fetal brain varies in US image volumes, initial global (affine) registration of the brain is expected to have a positive effect on the segmentation performance. Although the alignment of US volumes is a non-trivial task, methods have been developed for the automated alignment of fetal brain volumes [13, 14]. Furthermore, compared to the voxel-wise annotation of several subcortical structures, manual image alignment requires considerably less effort. For these reasons, alignment of the volumes to a common coordinate can be preferred over the additional voxel-wise manual annotation, and, as such, it is important to explore global alignment as a requisite preprocessing step.

Using few-shot learning, we aim to segment the choroid plexus (CP), lateral posterior ventricle horn (LPVH), cerebellum (CB) and cavum septum pellucidum et vergae (CSPV) during the second trimester of gestation. This trimester is of particular interest since this is when women undergo an US examination as part of routine care to screen for anomalies [15]. During this scan, several linear measurements, such as the transcerebellar diameter (TCD) and the atrial width, are collected and used for diagnostic purposes. Furthermore, in this trimester most subcortical structures have developed enough to be visible on US but are less affected by acoustic shading due to progressing calcification of the skull later in gestation. The specific subcortical structures studied in this work were selected based on their importance in this routine anomaly scan.

Lastly, to obtain an improved understanding of volumetric subcortical development during the second trimester, we will apply our developed segmentation models to a large cohort of healthy fetuses. The resulting model predictions will be validated for the CB, as volumetric growth curves for the CB have been reported in previous studies [16–22], and subsequently used to generate novel US-specific growth curves of subcortical volume development during the second trimester of gestation.

## 2. Related Work

Automated subcortical structure segmentation has been widely studied in MRI volumes of the adult brain [23–25]. However, the adult brain is structurally different to the actively developing fetal brain during gestation, which contains structures in transient stages of morphogenesis. For example, certain structures, such as the CSPV (which is in truth a fluid-filled cavity), are only present during gestation and typically disappear after birth [26]. As MRI can also be safely used to visualize the fetal brain, but is clinically only recommended to confirm or complement a diagnosis following an US examination [4], some studies have performed subcortical structure segmentation in fetal MRI volumes [27–31]. Notably, Gholipour et al. [30] published an annotated atlas of the fetal brain from 21 to 39 weeks of gestation and applied these labels to perform multi-atlas segmentation for new subjects. However, segmentation methods developed for MRI cannot be directly applied to US volumes due to the very different nature of the image acquisitions. While US is based on the interaction of the sound waves with tissue boundaries, resulting in discontinuous boundaries in the resulting image, MRI is based on protons in the tissue that react to magnetic fields. As a result, intensities on an MRI scan are rather uniform, whereas the intensities on an US scan heavily depend on the distance of the boundary to the probe as well as on the boundaries the US beam has already passed through. For example, the two lateral ventricles will appear very similar on a fetal brain MRI whereas on US typically only one lateral ventricle is well visible due to shadowing of the fetal skull in the proximal hemisphere. As a result of these acquisition differences, there is no direct one-to-one intensity mapping between US and MRI images, which makes it challenging to use fetal MRI atlas labels for US images.

In the remainder of this section we will outline previous work performed for subcortical structure segmentation in US. Furthermore, we will provide a brief overview of studies analyzing volumetric subcortical structure development during gestation.

### 2.1. Automated subcortical segmentation in US

Due to low soft-tissue contrast, high amount of speckle, and shadowing artifacts, automated anatomical segmentation in US volumes is a challenging task. Since most subcortical structures are small (i.e. the CB covers about 2.5% and the CP 0.8% of the intracranial volume at 22 GWs) and their boundaries are not always easily distinguishable, automated analysis of these structures has not been studied widely. Furthermore, manual segmentation is also challenging due to the aforementioned characteristics of US, and it can therefore be difficult to obtain accurate ground-truth labels. Before summarizing the past work, it is worth noting that the performance reported in different studies is not directly comparable due to the fact that the quality of US volumes can vary tremendously as a result of the transducer, acquisition protocol and scan settings. Additionally, performance measures such as the Dice Similarity Coefficient (DSC), which represent the percentage of overlap between ground-truth and prediction, are biased towards structural size, as for smaller structures the impact of a wrongly predicted voxel is much bigger than for bigger structures. For this reason, these should be carefully interpreted across the gestational age (GA) range, during which most structures grow substantially.

To the best of our knowledge, the first automated subcortical segmentation method for fetal brain US was developed by Gutiérrez-Becker et al. [32]. In that study, a statistical shape model was used to segment the CB in 3D fetal US volumes. The performance was promising, but it is not clear how this method can be easily extended to other subcortical structures with more shape variation than the CB, making it challenging to develop an initial shape model.

In another study [33], segmentation of the CP, LPVH, the CSP and the CB was performed in 3D US scans between 18 and 26 weeks of gestation. The authors proposed using a random decision forest using both appearance and distance features. Reported performance was good for the CP, LPVH and CSPV, but their method failed to generalize well to the CB. Furthermore, the search region for each structure was limited to the smallest cuboid enclosing all manual ground-truth annotations from the respective structure. More recently, Huang et al. [34] proposed using a region-based based descriptor to segment the CSPV (referred to as corpus callosum in the paper) and CP in 2D US. In contrast to the aforementioned studies, a 2D slice of the full fetal brain was used as input for the segmentation model. However, the sampled 2D slices were limited to standard US planes, making the segmentation substantially less challenging than full 3D segmentation.

Since the introduction of deep learning for image segmentation, most specifically the U-Net architecture [35], many segmentation tasks in medical imaging have shown an increase in performance. However, only a few studies have applied deep learning methods to subcortical structure segmentation in fetal US, as usually a relatively large dataset with ground-truth labels is needed. Obtaining voxel-wise labels for these structures, especially in 3D, is extremely time-consuming and challenging, due to the structural boundaries that are often hard to distinguish for a human annotator. One method that performs localization by predicting a bounding box, and included here for completeness, is the one developed by Huang et al. [36] that uses three 2D CNNs to localize several subcortical structures in 3D US (LPVH, thalami, CB and cisterna magna). Each of these networks predicted the orthogonal projections of the structures onto one of the three anatomical views (axial, sagittal and coronal), and the resulting 3D bounding box was then constructed from these 2D projections. However, this method is not suitable to extend to full voxel-wise segmentation in 3D as 2D projections can lack information needed to fully reconstruct an object in 3D.

Closely related to the study presented in this paper, Venturini et al. [37] used MRI atlas labels to overcome the need for manual annotation. Labels from the respective GW template of the fetal brain MR atlas [30] were registered to the individual US volumes, resulting in weak labels for the white matter, brainstem, thalami, and CB. These labels were subsequently used to train a multi-label 3D CNN to segment the respective subcortical structures in 3D US volumes. The reported performance was good, however, a severe downside of this study was that testing was performed on the same weak MR atlas labels as used in training. As the registered MR atlas labels can be erroneous, and do not always align well with the structures visible in the US volumes, this therefore yields a biased assessment of performance and as such does not analyze the effect of using weak labels. In Hesse and Namburete [9], active contours were applied to improve atlas-based labels as a pre-processing step before training a 3D U-Net, showing increased performance for CSP segmentation. However, that method was only validated for an age range between 20 and 24 GWs, and does not easily translate to all subcortical structures. Another method for segmenting the CSPV was presented by Wu et al. [38] and used a modified U-Net to perform segmentation in 2D US. However, as in Huang et al. [34], segmentation was limited to 2D standard planes.

In addition to the previously mentioned methods, some clinical studies report using multiplanar segmentation or Virtual Organ Computer-Aided AnaLysis (VOCAL) software (GE Healthcare) to perform their volume measurements in 3D US [17, 39–41]. These methods estimate the structural volume from 2D contours drawn on parallel (multiplanar) or oblique (VOCAL) planes. Although this does speed up analysis compared with fully annotating the volume, it still requires substantial manual annotation and is, as such, not considered to be an automated method.

### 2.2. Subcortical structure development

During a standard fetal US examination, multiple in-plane measurements, such as the TCD, are performed to assess the development and quantify the normality of the brain. For this reason, most studies analyzing subcortical growth during gestation use these 2D measurements [1]. Studies that do analyze volumetric development of subcortical structures during gestation most frequently use MRI volumes, either by manually annotating the volumes [19, 20, 22] or by using an atlas-based approach [42]. Theoretically, volumetric measures should be independent of the acquisition type (MRI or US), but in practice these measures can vary due to the different appearance of tissues in each modality. Furthermore, subcortical structures that are easily visible on US, such as the CP, are much harder to discern on fetal MRI. For these reasons, US-specific growth curves are needed.

A small number of studies have analyzed 3D subcortical development using US [17, 18, 21, 39–41, 43, 44], and all have used the previously described multiplanar or VOCAL software. In Benavides-Serralde et al. [40], total intracranial, frontal, thalamic and cerebellar volumes were measured between 28 and 34 weeks of gestation and these measurements where used to estimate the average volume differences between a group of 39 growth-restricted fetuses and a group of 39 age-matched appropriate-for-gestational age fetuses. The same structures were measured between 20 and 36 GWs and used to compare the structural volumes of 73 fetuses with congenital heart disease to a group of 168 normal control fetuses [39].

In order to construct normal ranges for thalamic volume development during gestation, Sotiriadis et al. [41] measured the thalamic volume of 122 normal fetuses between 20 and 34 weeks. Furthermore, it was shown that there were no significant volumes differences between the left and right thalamus. Setting out to generate reference values and growth curves for 3D US, Babucci et al. [17] obtained volumetric measures for the thalamus, CB and cortex for 344 fetuses from 22 to 39 weeks. Only a small number of volume measurements were, however, included after 30 weeks, due to shadowing of the fetal skull. Focussing only on cerebellar growth, multiple studies have developed US-specific growth curves for this structure from a healthy population [18, 21, 43, 44].

In summary, even though several studies have proposed subcortical segmentation methods for fetal US, a single method obtaining good performance across multiple structures is lacking for 3D US. Furthermore, due to the absence of accurate automated segmentation methods, only thalamic and cerebellar volumetric development during gestation have been studied in previous work. In this study, we address these limitations by developing a segmentation method aiming to obtain competitive segmentation performance across multiple subcortical structures (CB, LPVH, CSPV and CP) and subsequently applying these methods to obtain novel volumetric growth curves for the respective structures during gestation.

## 3. Methods

In this study, we aim to accurately segment the CB, LPVH, CSPV and CP in 3D US image volumes during the second trimester of gestation using a minimal number of voxel-wise annotations, i.e. few-shot segmentation. Defining *n_a_* as the number of images that is manually annotated, we will consider two approaches, both obtained from an equivalent number of manual annotation: (1) training a model naively with *n_a_* individually annotated 3D images, and (2) training a model with a large number of weak propagated atlas labels obtained from annotating *n_a_* 3D template images.

In this section, we start by introducing some notation, followed by an explanation of the network architecture used for training. Next, the fewshot training labels are explained in more detail, including both the template image generation and the manual annotation protocol. In the experimental section, an outline will be given of the data used in this study as well as the experiments performed and the evaluation metrics. Finally, in the last section we will present how the subcortical growth curves are generated.

### 3.1. Notation

To facilitate the methods presented in this section, we will introduce some notation. Given a dataset of *m* image volumes, the set of image volumes in their original acquired orientation will be denoted as 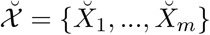 with 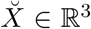 and the set of image volumes rigidly aligned to the same reference coordinate system as 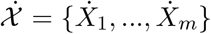. Referring to either 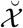 or 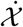 will simply be done by using ***χ***. Furthermore, we will refer to the original and manually aligned images as *unaligned* and *aligned*, respectively.

Multi-label segmentation masks will be denoted by *Y* . Referring to the binary mask of a single class will be done by the subscript *c*, with *c* being the class (structure), and the type of the label (described in next section) will be marked with a superscript 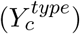. Model predictions are marked with a hat over them, and follow the same convention to denote the class 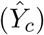. As introduced before, *n_a_* refers to the number of manually annotated (template) images used for training in a certain experiment. A full overview of all notation used in this work can be found in Appendix Appendix A.

To avoid ambiguity between *image volumes* and a *volume measurement*, in the remainder of this paper we will refer to our 3D image volumes simply as *images*.

### 3.2. CNN architecture

We propose a multi-label 3D U-Net [35] with batch normalization [45] for the subcortical segmentation. The network, defined by *θ*, predicts a multi-channel segmentation for a 3D US input image: 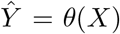. Following the same convention as for the images, 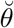 and 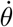 correspond to networks trained with 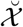 and 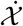 respectively.

The network predicts an output prediction with *k* channels, in which *k* is equal to the number of segmented classes. To obtain probabilities for each individual class, a soft-max function is applied across the channel dimension of 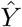 . Final predictions for a single class, denoted by 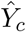, are subsequently obtained by assigning each voxel to the class with the highest probability. A schematic overview of the segmentation network is shown in Fig. 1.

**Figure 1:**
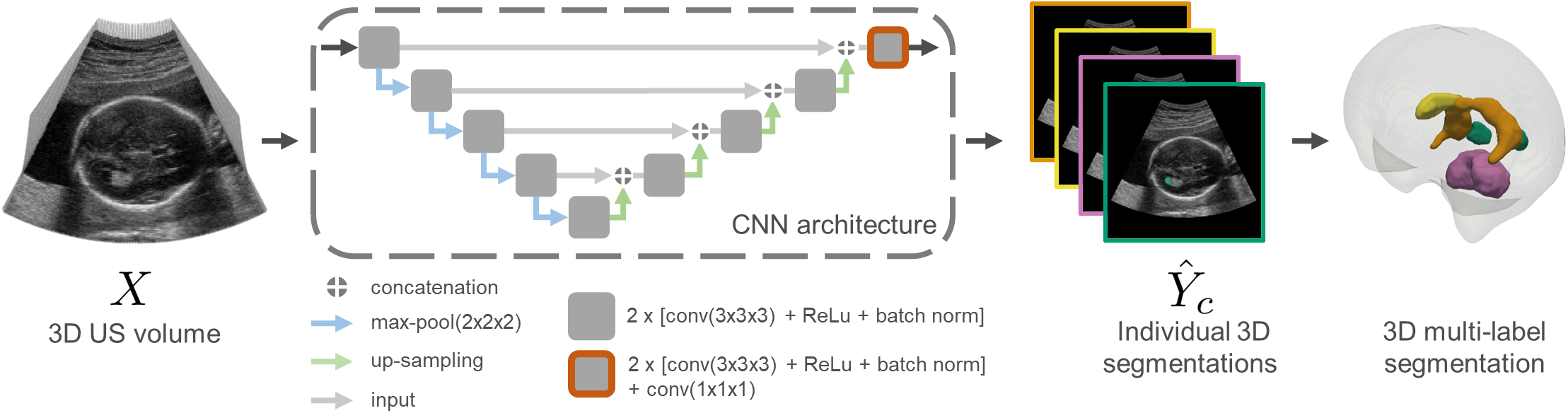
Schematic overview of segmentation network. The segmented structures shown are the CB (pink), LPVH (green), CSPV (yellow) and CP (orange). The LPVH and CP are shown in both hemispheres for visualization purposes, but are only segmented in the visible hemisphere.

The U-Net architecture consists of two symmetrical branches, an encoder and a decoder, linked by skip connections. Both the encoder and decoder are composed of several convolutional blocks, each containing two 3D convolutional layers (kernel size of 3×3×3), followed by a ReLU activation function and batch normalization [45]. In the encoder branch, each block is followed by a max-pool layer performing down-sampling, whereas in the decoder branch each block is followed by an up-sampling operation. The skip connections link the output of each block in the encoder to the corresponding block in the decoder branch. To reduce the number of channels in the output to the number of classes, the last block of the decoder branch contains an additional convolutional layer of kernel size 1×1×1.

In this study, we set *k* to five, resulting in predictions 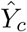 with *c* ∈ {*CP, LPV H, CSPV, CB, background*}. Based on initial experiments, we designed our network to have a depth of five, defined as the number of blocks in the encoder pathway and empirically set the number of feature maps in the convolutional layers to 16 in the first block of the encoder. For each subsequent block, the number of feature maps was multiplied by two in the encoder branch, and divided by two in the decoder branch. In the decoder pathway, we used nearest-neighbor interpolation for the upsampling operation. However, although we obtained best performance with this network configuration during initial experiments, we are in this study not particularly interested in the effect of the network architecture on the segmentation task but aim to explore the optimal way of training a CNN with a small number of manual annotations.

### 3.3. Few-Shot Training Labels

As described before, we consider two types of ground-truth labels (1) a small set of expert annotations, obtained from individually annotated images, and (2) a large set of weak labels obtained from propagated atlas annotations. These will be referred to as *expert* (*Y^exp^*) and *atlas* (*Y^atl^*) labels respectively. Likewise, models trained with either expert or atlas labels are denoted by *θ^atl^* and *θ^exp^*. In the next sections, we describe the generation of the atlas labels as well as the manual annotation process in more detail.

#### 3.3.1. Atlas Labels

Atlas-based labels were obtained for the images by manually annotating 3D template images as opposed to annotating individual images (Fig. 2). Template images were created by registering a subset of *s* rigidly aligned images 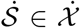 to each other using a Demons diffeomorphic groupwise registration approach [46], resulting in a set of non-rigid transformations that can transform each image from the rigidly aligned orientation to a reference space *R* defined by: 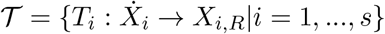. The registered images were subsequently averaged to obtain a template image 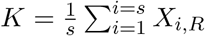.

**Figure 2:**
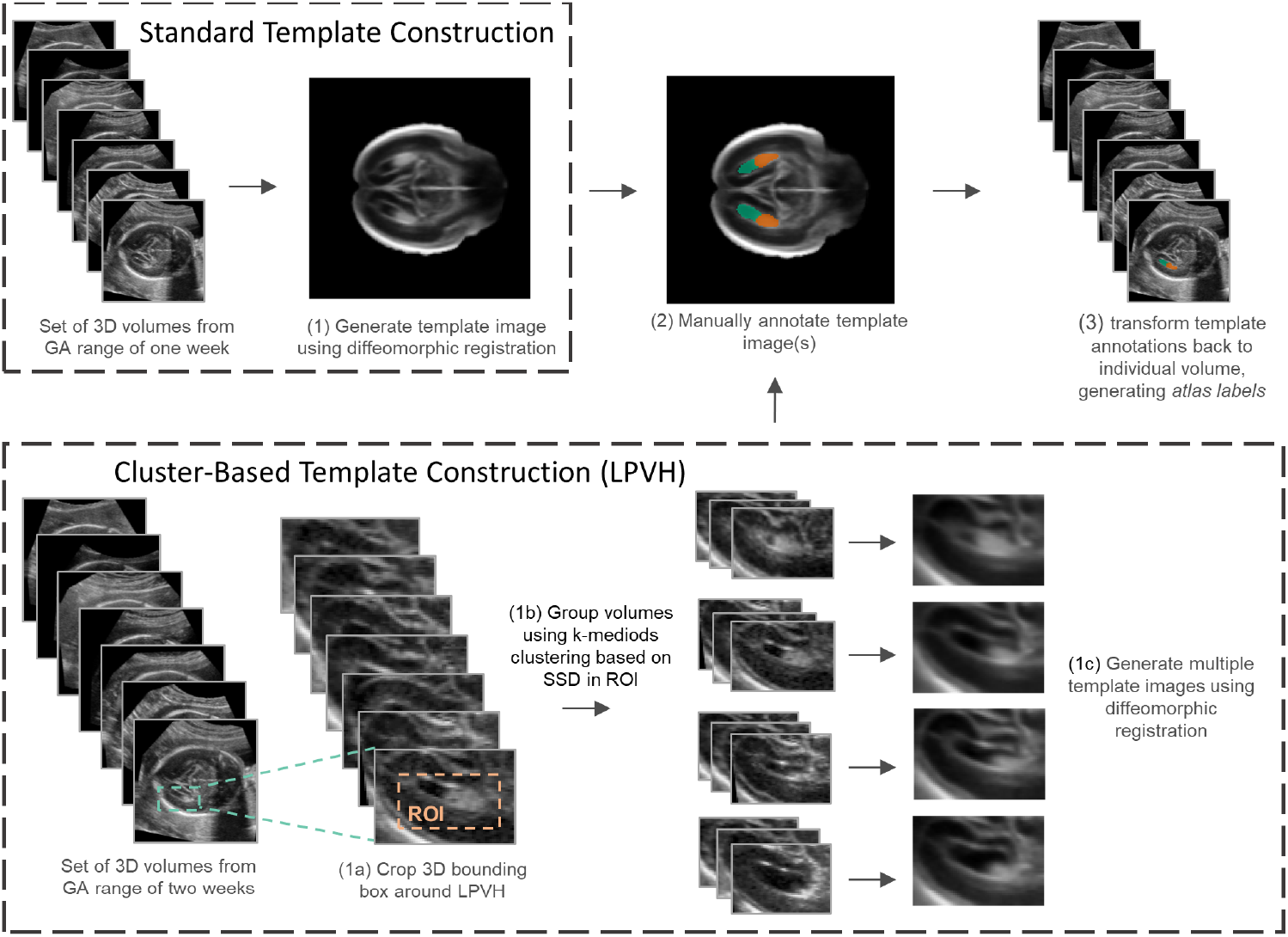
Schematic overview of atlas label generation. Standard whole brain templates (top row) were constructed for each GW and all structures (CB, CP, CSPV and LPVH) were annotated in these templates. Cluster-based template construction (bottom row) was only performed for the LPVH.

In this template *K*, the four subcortical structures of interest were manually annotated (see Section 3.4), resulting in a set of labelled template images. For simplicity, the annotated templates images will also be referred to with *K*. Subsequently, these labels were non-linearly registered back to the individual images using the inverse transform of the registration to the reference space, given by: 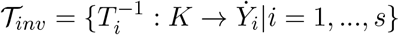. Due to anatomical variations between individuals and potential errors in the registration, the resulting labels may contain imperfections and are thus considered to be weak labels. To clearly differentiate between the annotated template images (which could be referred to as an atlas) and the propagated atlas-based labels, we will refer to the former only as *annotated template images* and to the latter as *atlas labels*.

##### Standard Template Construction

In order to generate the template images, rigidly aligned images from a GA range of 1 week were used, resulting in one template image per GW. To capture possible differences in anatomy between the left and right cerebral hemispheres, template images for each hemisphere were constructed separately using the respective images and fused together at the midsagittal plane by joining the interhemispheric fissure in the two templates (Fig. 2 top row). The set of annotated template images (one per GW) is denoted by 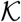, containing annotations of all segmented structures (LPVH, CB, CP and CSPV).

##### Cluster-Based Template Construction

During initial experiments with the atlas labels, it was seen that the LPVH shape was not well captured by a single template image per GW. For this reason, our method was extended to generate multiple templates for this structure using a clustering approach (Fig. 2 bottom row). We selected images from an age range of two GWs, grouped them using k-mediods clustering and subsequently constructed four template images of the LPVH using this grouping. As any anatomical variation between left and right can be captured by the clustering, images of the left and right hemisphere were combined by flipping the right side images across the midline. In the resulting clustered template images (denoted by *K^clust^*) only the LPVH was manually annotated, resulting in an additional set of atlas labels for the LPVH. Unless explicitly mentioned otherwise, the labels propagated from the cluster-based template images are used for the LPVH in the remainder of this study (i.e. 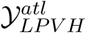 is obtained from the cluster-based template annotation).

To perform the k-mediods clustering, all images were cropped to an area around the LPVH (same bounding box dimensions across the whole GA range) and the intensities of the cropped images were normalized using histogram equalization. The size of this crop was empirically set, and was pre-dominantly performed to reduce computation time. Next, all bounding boxes were pairwise registered to each other (using the same Demons registration used for template construction) and the sum-of-squared-differences (SSD) was computed over a region of interest (ROI) around the LPVH. This ROI was manually set for every two-week gestational window. The resulting square matrix of pairwise SSD values was subsequently used as input for the k-mediods clustering. As k-mediods clustering can be sensitive to outliers, we excluded a few outliers based on their visual appearance prior to clustering. After the clustering step, each outlier was assigned to the nearest cluster based on the distance to the cluster center. Once all images were assigned to a cluster, templates were created from the images in each cluster, including the outliers, as described earlier. The k-mediods clustering was set to repeat 500 times, and results containing clusters with less than three images were discarded.

### 3.4. Manual Annotation Process

Both the constructed template images, 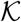 for all structures and 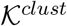 for the LPVH, as well as a subset of individual images, 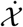, had to be manually annotated. Since subcortical segmentation is not a task usually performed clinically, and structural boundaries can be very challenging to distinguish in 3D US images, a bespoke segmentation protocol was defined in consultation with experienced ultrasonographers (M. Aliasi and M.C. Haak). For each structure, the 2D views in which the structural boundaries were best visible were identified (see Table 1). Based on this protocol, the manual segmentations (for both template and individual images) were performed by L.S. Hesse and verified by M. Aliasi. To illustrate the challenging nature of subcortical structural segmentation, an example annotation of the CB across the three orthogonal views can be found in the Appendix (Fig. B.12).

**Table 1:**
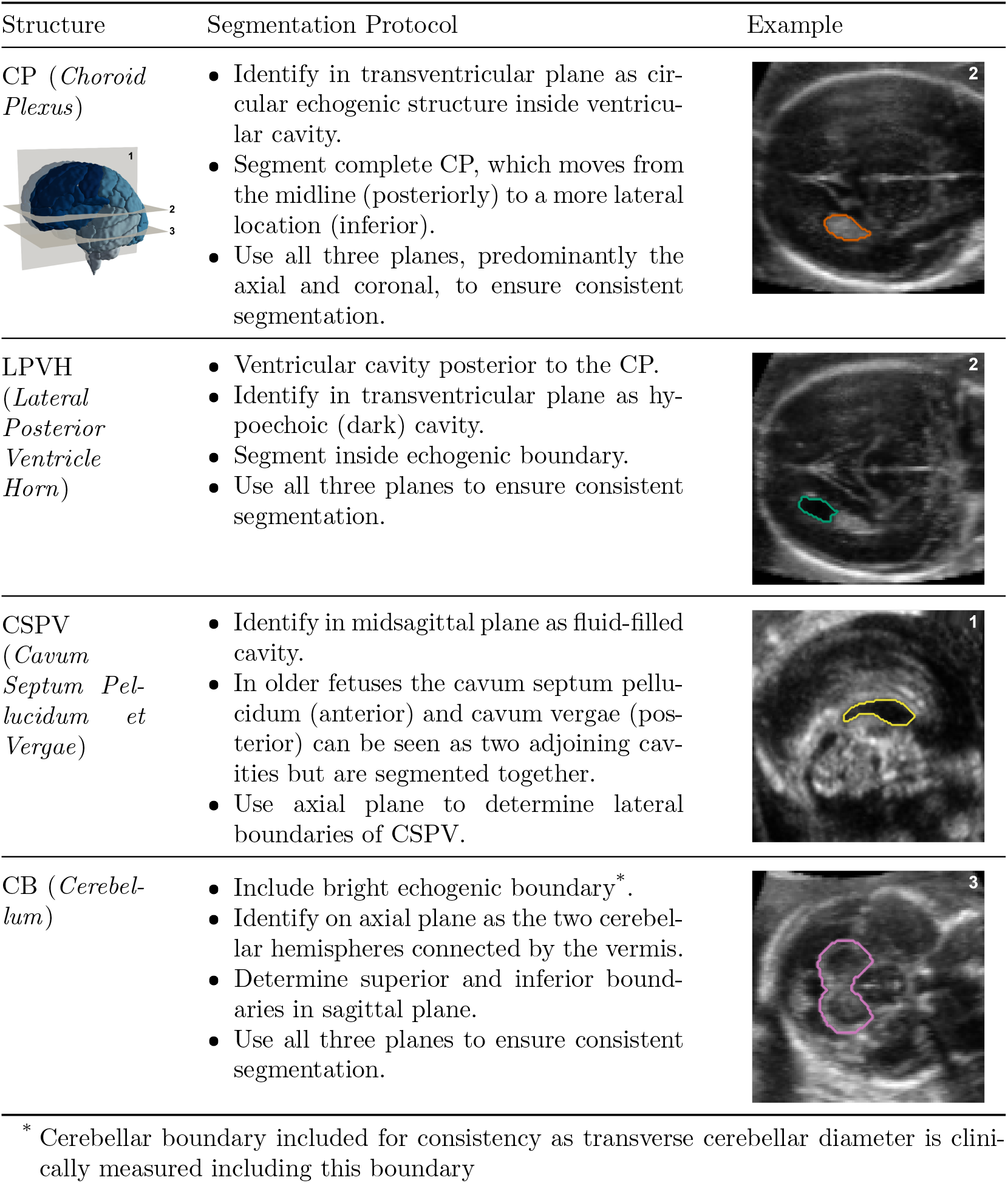
Overview of manual segmentation protocol

| Structure | Segmentation Protocol | Example |
| --- | --- | --- |
| CP (*Choroid Plexus*) | Identify in transventricular plane as circular echogenic structure inside ventricular cavity.<br>Segment complete CP, which moves from the midline (posteriorly) to a more lateral location (inferior).<br>Use all three planes, predominantly the axial and coronal, to ensure consistent segmentation. |  |
| LPVH (*Lateral Posterior Ventricle Horn*) | Ventricular cavity posterior to the CP.<br>Identify in transventricular plane as hypoechoic (dark) cavity.<br>Segment inside echogenic boundary.<br>Use all three planes to ensure consistent segmentation. |  |
| CSPV (*Cavum Septum Pellucidum et Vergae*) | Identify in midsagittal plane as fluid-filled cavity.<br>In older fetuses the cavum septum pellucidum (anterior) and cavum vergae (posterior) can be seen as two adjoining cavities but are segmented together.<br>Use axial plane to determine lateral boundaries of CSPV. |  |
| CB (*Cerebellum*) | Include bright echogenic boundary*.<br>Identify on axial plane as the two cerebellar hemispheres connected by the vermis.<br>Determine superior and inferior boundaries in sagittal plane.<br>Use all three planes to ensure consistent segmentation. |  |

\* Cerebellar boundary included for consistency as transverse cerebellar diameter is clinically measured including this boundary

As described before, typically only one hemisphere is well visible in the 3D US images. For this reason, annotation of the CP and LPVH in 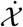 was only performed in the most visible hemisphere. However, as the whole brain template images 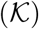 were created for both the left and right hemisphere separately and subsequently fused together, manual annotation of these images involved annotation of the CP and LPVH in both hemispheres, thus resulting in two manual template annotations per GW for these structures. As both the CSPV and CB are structures near the midsagittal plane, these are less affected by acoustic shadowing and are segmented in full. As a result, when setting an equal *n_a_*, defined as the number of manually annotated (template) images, between expert and atlas labels, this results in an equivalent number of annotations for the CB and CSPV, but twice as much annotation of the CP and LPVH for the atlas labels.

The annotation of the LPVH in both left and right hemisphere in the standard template images also means that the annotation effort required to annotate the LPVH in 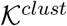 (four templates per two GWs) was the same as annotating the LPVH in 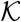 (one template per GW, annotating both the left and right LPVH). For consistency among the structures, for the atlas labels *n_a_* will refer to the number of standard template images that were annotated, which is thus equal to one per GW.

All manual annotations were performed on the aligned images, 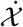 . Labels for 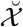 images were generated by transforming the labels back to the unaligned space using the inverse of the rigid manual alignment transform.

### 3.5. Experimental Set-Up

#### 3.5.1. Dataset and Preprocessing

For this study, 3D US images from the INTERGROWTH-21^st^ Fetal Growth Longitudinal Study have been used [47]. The main aim of that project was to study growth, health, nutrition and neurodevelopment in a multi-site ethnically diverse population. Based on a list of strict criteria such as, age, BMI and medical history, healthy women were included from eight carefully selected sites around the world. All images were acquired from the axial plane on a Philips US machine (Philips HD-9, Philips Ultrasound, USA) with a curvilinear abdominal transducer. We used a total of 537 images of which the fetuses were born without congenital malformations, between the age range of 18 and 26 GWs, inclusive. Furthermore, it was ensured that images from both the left and right hemisphere were included and that only images with sufficient US quality were included in this total. In Fig. 3 the data distribution over the GA range is shown.

**Figure 3:**
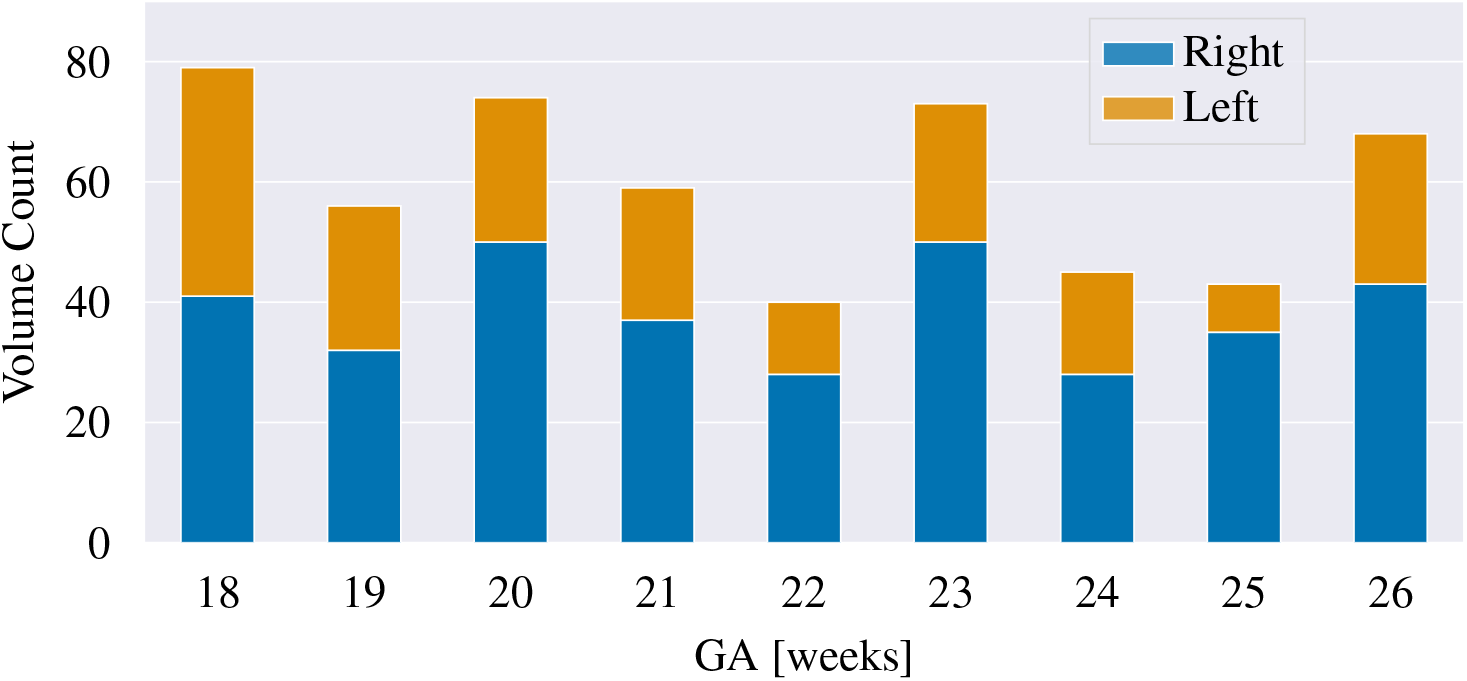
Data distribution of the 537 US images used in this study

As an initial pre-processing step, all images were resampled to an isotropic voxel size of 0.6 mm (using trilinear interpolation) and cropped to the same dimensions of 160 × 160 × 160 voxels. Furthermore, all images were manually aligned using a rigid transformation to the same reference space, as the anatomical orientation of the brain varies in the original scan due to transducer and fetal position during acquisition.

The complete dataset consisted of 537 images, of which 259 were used for model development and the remaining 278 to generate volumetric growth curves of the four subcortical structures (referred to as the *analysis* subset or ***χ***_*analysis*_). Of the 259 images used for model development, 20 were used for testing (***χ***_*test*_) and the remaining for training (***χ***_*train*_) and validation (***χ***_*val*_). These 20 test set images were evenly distributed across the GA range that we used, with four images for every second GW (at 18, 20, 22, 26, and 26 GWs). Furthermore, it was ensured that the training, validation, and analysis subsets were also evenly distributed across the gestational range. All testing images were manually annotated and thus had expert ground-truth labels (as described in 3.4). Ten test images were twice manually annotated by the same observer, in order to obtain intra-observer variability.

To obtain atlas labels for our dataset 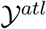, template images had to be generated for our dataset (*K* and *K_clust_*). These template images were created using all images in our dataset (***χ***_*train*_,***χ***_*val*_,***χ***_*test*_, ***χ***_*analysis*_), as these were available from earlier work [46]. However, the effect on the appearance of the template images of using all images versus only the one used for training is very small, and as such not expected to have a significant effect on the resulting atlas labels.

As the GA of the dataset used in this study ranges between 18 and 26, *n_a_* for the training data was set to 9. This yielded two overlapping training sets, one with 239 images containing weak atlas labels, and one with 9 images containing expert labels. During training of our networks, 10% of the 239 was used as validation. For the 9 expertly labelled individual images, no validation set was used, but all settings were kept the same as when training with atlas labels. An overview of how the data were separated into different subsets, and the number of available labels in each set is summarized in Table 2.

**Table 2:** Overview of the number of images and available labels for each of the data subsets

|  | volumes | atlas labels | expert labels |
| --- | --- | --- | --- |
| Analysis subset | 278 | 0 | 0 |
| Model Development subset |  |  |  |
| Testing | 20 | 20 | 20 |
| Training | 215 | 215 | 9 |
| Validation | 24 | 24 | 0 |

#### 3.5.2. Experiments

In order to study the effect of using only a small number of manual annotations, multiple experiments were performed.

##### Comparison between atlas and expert labels

Firstly, we aimed to study the effect of using *n_a_* expertly annotated images versus many weakly labelled images, obtained from annotating *n_a_* template images. This first experiment was performed with images from the full GA range in our data, resulting in *n_a_* = 9. Defining *n_x_* as the number of training images used during training, two separate networks were trained: one that used all atlas-labelled images (*θ^atl^*, *n_x_* = 215) and a second that only used the expertly labelled images (*θ^exp^*, *n_x_* = 9). As the alignment of the images was expected to influence the performance, this experiment was repeated for the aligned images, 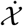, as well as for images in their original orientation, 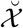. Models trained with aligned images were evaluated on aligned testing data, and vice-versa.

##### Varying the number of manual annotations

Next, *n_a_* was reduced to study the performance decrease when using even fewer manual annotations for training. For 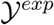, a simple subset of *n_a_* images were used whereas for 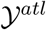 all images obtained from *n_a_* annotated templates were used. The models trained with these labels are noted as 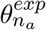 and 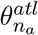, and in our experiments *n_a_* ranged from 2 to 9. The images used for training were selected to be uniformly distributed over the whole GA range, e.g. for *n_a_* = 2 the images from 20 and 24 GWs were selected. Even though these models were only trained with images from selected GWs, evaluation was performed on all testing images. As for the previous experiment, this was repeated for both 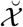 and 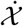 .

##### Split training by GA

We then split our training data by GA and trained separate networks for each two-week gestational window. Testing was sub-sequently only performed on the testing images of the corresponding GWs. These models trained per two-week window are referred to as 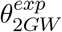 and 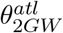 and are compared to the networks trained on the full GA range, *θ^exp^* and *θ^atl^*. The aim of this experiment was to study the effect of combining the whole GA range in a single model versus training models for smaller gestational windows. The images in the dataset used span an uneven ges-tational range of 9 weeks (from 18 to 26 GWs). For this reason, as 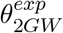 is trained using a two-week gestational window, no network was trained with the images of 26 GWs. As the effect of alignment is explored in the first two experiments, we only performed this experiment using 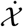 .

##### Cluster-based atlas labels

During initial testing, we observed that using atlas labels from multiple LPVH templates, *K^clust^*, increased performance compared to using the labels from a single annotated template per GW, *K*. For this reason, all aforementioned experiments using atlas labels were performed with the LPVH annotations from the clustered template images. To quantify the performance increase from these improved template images, we also annotated the LPVH in *K*, and trained a network with these labels. As this only applies to the atlas labels, no expert training labels were used in this experiment.

#### 3.5.3. Network Training

To train our networks, we used a combination of multi-class DSC 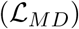 and Cross-Entropy (CE) loss 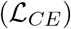, as defined by:

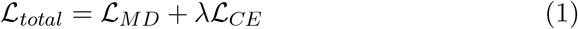

with *λ* as a relative weighting parameter between both loss terms. It was shown during initial experiments that the DSC loss term resulted in higher performance for individual structures due to the high unbalance between the structures and background, whereas the addition of the CE loss term ensured convergence for all structures. The *λ* was set to 1 based on initial experiments, but did not have a strong effect on performance for values within the same order of magnitude.

To prevent overfitting, for each sample a combination of the following geometric augmentations was performed: horizontal flips (across the midline for aligned images), rotation (±30°), translation (±10 voxels) and scaling. The scaling range (defined by *s_min_* and *s_max_*) was set for each GW separately and given by:

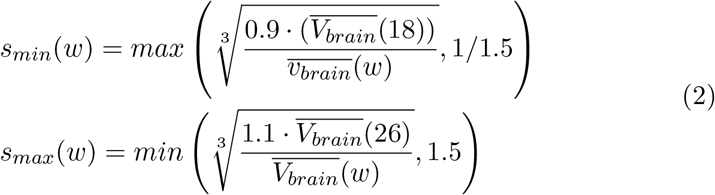

with *w* the GA of the fetus in weeks and 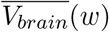 the average whole brain volume at a certain GW. This ensured that scaling was only performed to brain sizes within the dataset (i.e. images at 18 GW were predominantly up-scaled whereas images at 26 GW were mostly down-scaled). We applied augmentation for training with both 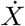 and 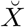 . For 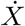, this meant that the network was trained with images slightly deviating from the initial alignment to the same coordinate system, making it more robust to imperfect alignment.

As the size of the training set varies strongly across experiments, all our models were trained for the same number of iterations (defined as passing a single batch of data through the network). This number was empirically set to be equivalent to training with all atlas-labelled images for 100 epochs. We chose the ADAM optimizer [48] with an initial learning rate of 0.001 for training, and used a batch size of four. This relatively small batch size was constrained by the GPU memory due to the large input images (160 × 160 × 160). The results reported in this study are the performance after the last training epoch averaged over three training runs.

#### 3.5.4. Implementation

All models were implemented in Python 3.7 using PyTorch (version 1.7.1). We trained our models on an NVIDIA Tesla V100 with 32 GB of memory. To generate the template images used to obtain our atlas labels we used Matlab (version 9.8), and all manual annotation was performed using the freely available MITK-Workbench.

### 3.6. Evaluation

#### 3.6.1. Postprocessing

During visual inspection of the resulting segmentations, we noticed that for some images (about 25% of the test images for 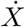 and 50% for 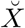), *θ^exp^* made predictions that were also in the hemisphere proximal to the transducer (which is partly occluded due to shadowing), whereas our ground-truth only contained manual annotations of the distal hemisphere for both the CP and LPVH. However, these predictions were always smaller in size than the predictions in the distal hemisphere, as only part of the structure was clearly visible. Furthermore, for 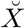 some very small spurious areas were predicted for a few test images. For these reasons, we post-processed our predictions 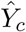 by taking the largest connected component as the final prediction, denoted by 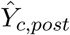. As the evaluation metrics are computed both with and without post-processing, for simplicity the notation 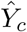 is used in the remainder of this section to represent either case.

#### 3.6.2. Metrics

The predictions for each class were separately evaluated using the following metrics, where 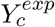 is a binary manual expert label in the test set for a single class, and 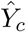 the predicted binary mask for this single class:

- The Dice Similarity Coefficient (DSC), defined by:

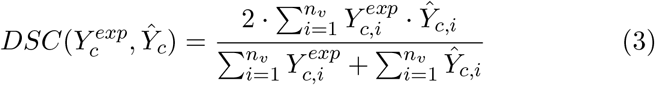

with *n_υ_* the total number of voxels per image. A DSC of 1 indicates perfect overlap between the ground-truth and prediction, whereas a DSC of 0 indicates no overlap.
- The 95^*th*^ percentile Hausdorff distance (*H*_95_), which is the maximum (or 95^*th*^ percentile in our case) over the set of boundary points of the minimum distances (between a boundary point in one set and any boundary point in the other set). If *B_c_* and 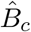 represent the surface points of *Y_c_* and 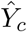 respectively, the *H*_95_ is given by [49]:

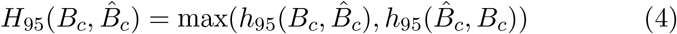

where *h* is the directional Hausdorff distance given by:

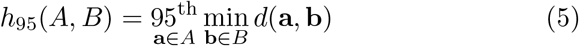

with *d*(**a**, **b**) the Euclidean distance between points **a** and **b**, and 95^*th*^ being the 95^*th*^ percentile. We are using the 95^*th*^ percentile distance because it is more robust to very small outliers than the standard maximum Hausdorff distance. As the Hausdorff distance represents the distance between prediction and ground-truth, lower distances indicate better segmentation performance.
- The signed (Δ*V_rel_*) and unsigned (|Δ*V_rel_*|) relative volume differences, defined by:

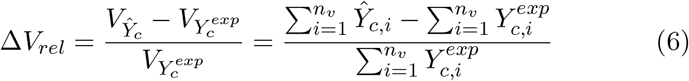

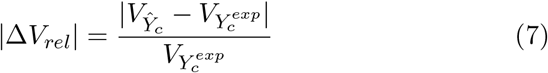

The signed difference indicates whether the model’s prediction is over- or under-segmented. A positive number indicates over-segmentation, whereas a negative number indicates under-segmentation. On the other hand, the unsigned volume difference indicates the error in the volumetric measures predicted by the segmentation network. Given that the segmentation networks were developed in order to extract volumetric information of the structures, this measure is important in estimating the expected error. As the structures segmented in this study vary considerably in size, the relative volume differences (with respect to the ground-truth structural volume) enable comparison across the structures.

### 3.7. Volumetric Subcortical Growth Trajectories

In order to generate growth curves for the subcortical structures, we finally applied our models to the US images in our analysis subset. We predicted subcortical volumes 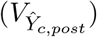 for 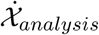 using both 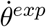 and 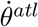 in order to analyze the differences. Subsequently, a linear or quadratic polynomial was fit to the predicted volumes as function of the GA. The quadratic term was only added if it proved to be statistically significant for both 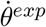 and 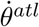 (determined by a two-sided t-test), as the underlying structure growth should be the same between the two networks.

In addition to structural volumes, we also computed the relative structural volume with respect to the whole brain volume, defined by:

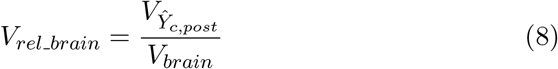

with *V_brain_* the whole brain volume of the respective image. These whole brain volumes were computed from whole brain masks derived from an MRI fetal brain atlas [30]. The template images in this atlas, ranging from 21 to 31 GWs, where first binarized to obtain whole brain atlas masks and subsequently aligned to the individual US images using an affine transform (rigid + scale). For each US image, the template of the corresponding GW was used, and for fetuses younger than 21 GWs, the template image of 21 GWs was used.

### 3.8. Ethics Statement

The INTERGROWTH-21^st^ Project was approved by the Oxfordshire Research Ethics Committee “C” (ref: 08/H0606/ 139), the research ethics committee of the individual participating institutions and the corresponding regional health authorities in which the project was implemented. Participants provided written consent to be involved in the project [50].

## 4. Results

All performance values presented in this section were obtained from 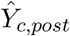 (predictions post-processed to only keep the largest connected component). Results without post-processing are given in the Appendix (Figs. B.13 and B.14).

### 4.1. Weak Atlas Labels

In order to quantify the amount of inaccuracies in our weak atlas training labels, we determined the DSC overlap between the naive atlas propagated labels 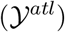 and the manual ground-truth labels 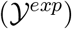 in our test set. These results are shown in Table 3 as *prop. atlas*. For the LPVH, the DSC overlap of the atlas labels obtained from the clustered template images, *K^clust^*, is shown. It can be observed that the naive propagated atlas labels obtained relatively low DSC, ranging from 0.68 for the LPVH to 0.80 for the CB.

**Table 3:**
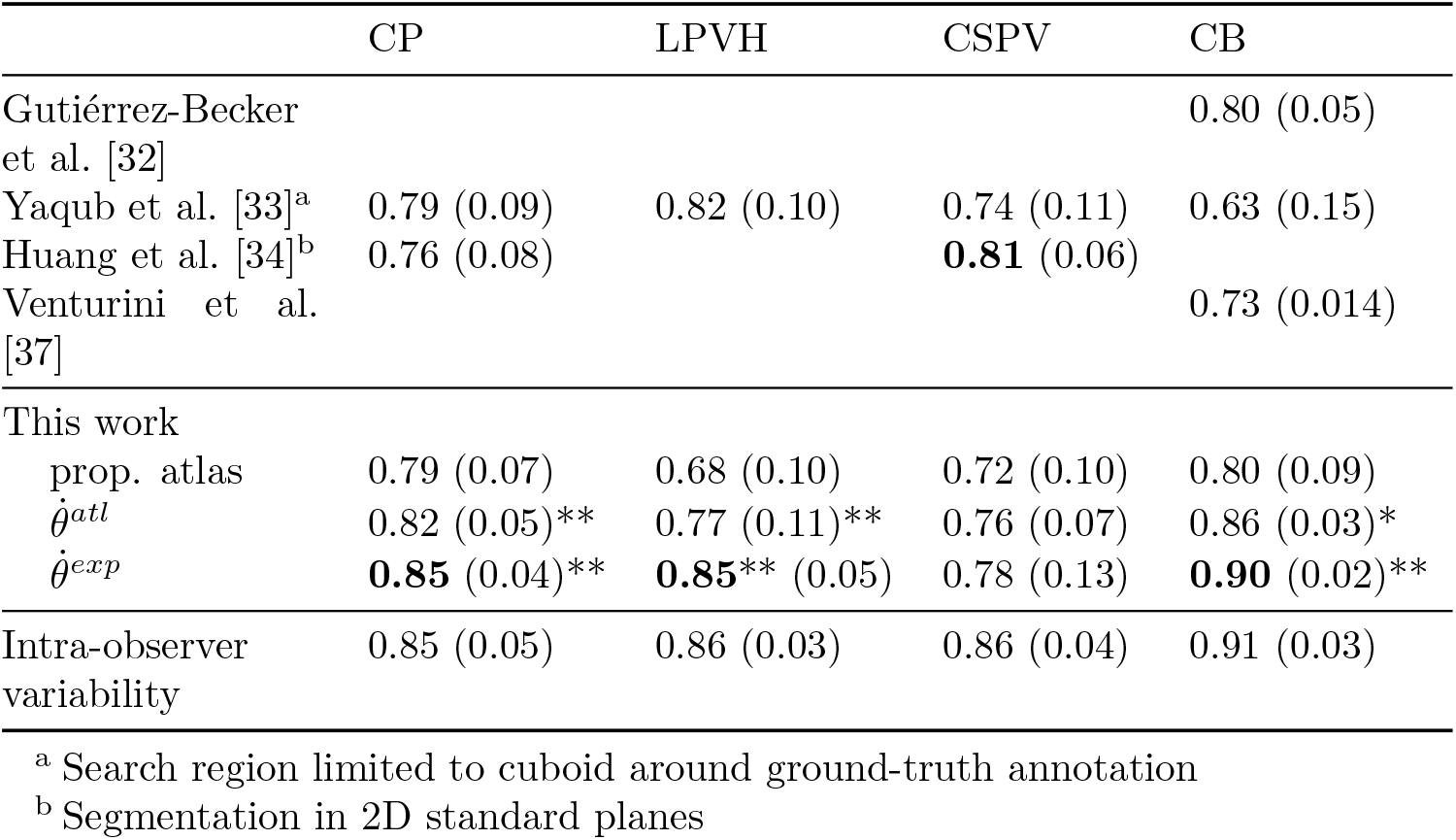
DSC performance of our segmentation networks using aligned images compared to results of previous studies (obtained on different datasets). Numbers in between brackets indicate the standard deviation and the best performance for each structure is shown in bold. For both 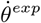 and 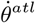 results of statistical testing with respect to the propagated atlas labels are shown with (**) *p* < 0.005 and (*) *p* < 0.05. Full overview of statistical results can be found in Appendix C.

### 4.2. Comparison between atlas and expert labels

#### Effect of image alignment

The resulting segmentation performance of our trained networks *θ^atl^* (*n_x_* = 239) and *θ^exp^* (*n_x_* = 9) are shown in Fig. 4 and Table 3. The results shown were based on experiments that were trained and tested on images with the same alignment, either 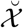 or 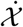. Table 3 shows the performance for the aligned images whereas in Fig. 4 results for both the aligned and unaligned settings are presented. From Fig. 4 it can be seen that higher performance is obtained for the aligned images compared to the unaligned images, which is most pronounced in the lower *H*_95_ for these images. For the aligned images, 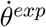 outperforms 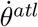 in terms of DSC for all structures. This trend is also seen when comparing the *H*_95_, except for the CP. On the other hand, on the unaligned images both 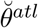 and 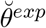 obtain similar performance across the structures.

**Figure 4:**
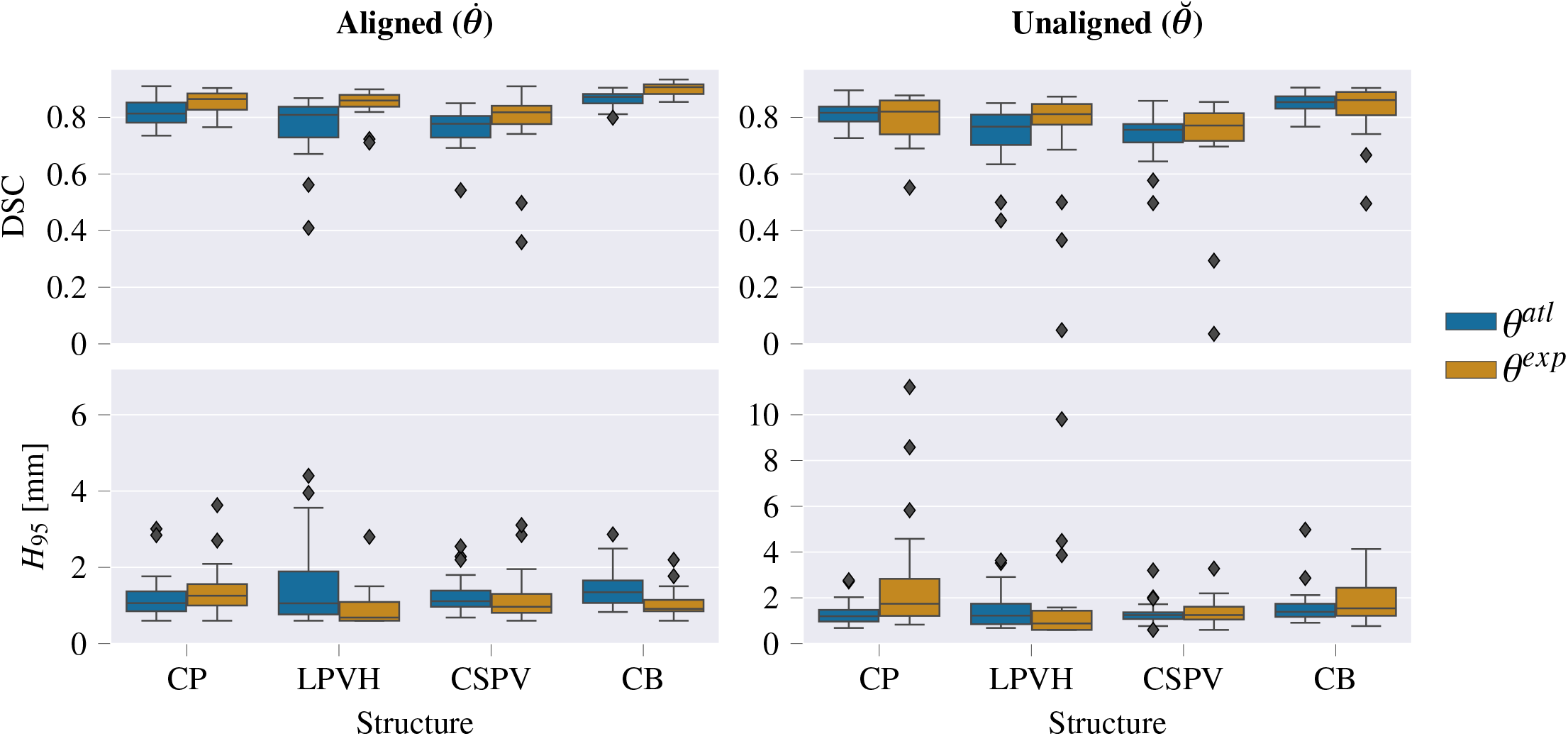
Resulting performance values after post-processing for *θ^exp^* and *θ^atl^*.

#### Volumetric measures

In Fig. 5 the resulting relative volume differences (Δ*V_rel_* and |Δ*V_rel_*|) are shown. It can be observed that *θ^atl^* shows an average Δ*V_rel_* higher than 0 (except for the CSPV in the aligned setting), corresponding to over-segmentation. In contrast, *θ^exp^* shows, on average, a slight negative Δ*V_rel_*, thus under-segmentation, for the unaligned images while the differences are centered around zero for the aligned data. In Fig. 5 also some clear outliers for both the CSPV and LPVH can be observed. However, these correspond to images at 18 or 20 GW where the total ground-truth volume of the respective structures is very small, resulting in large relative volume differences.

**Figure 5:**
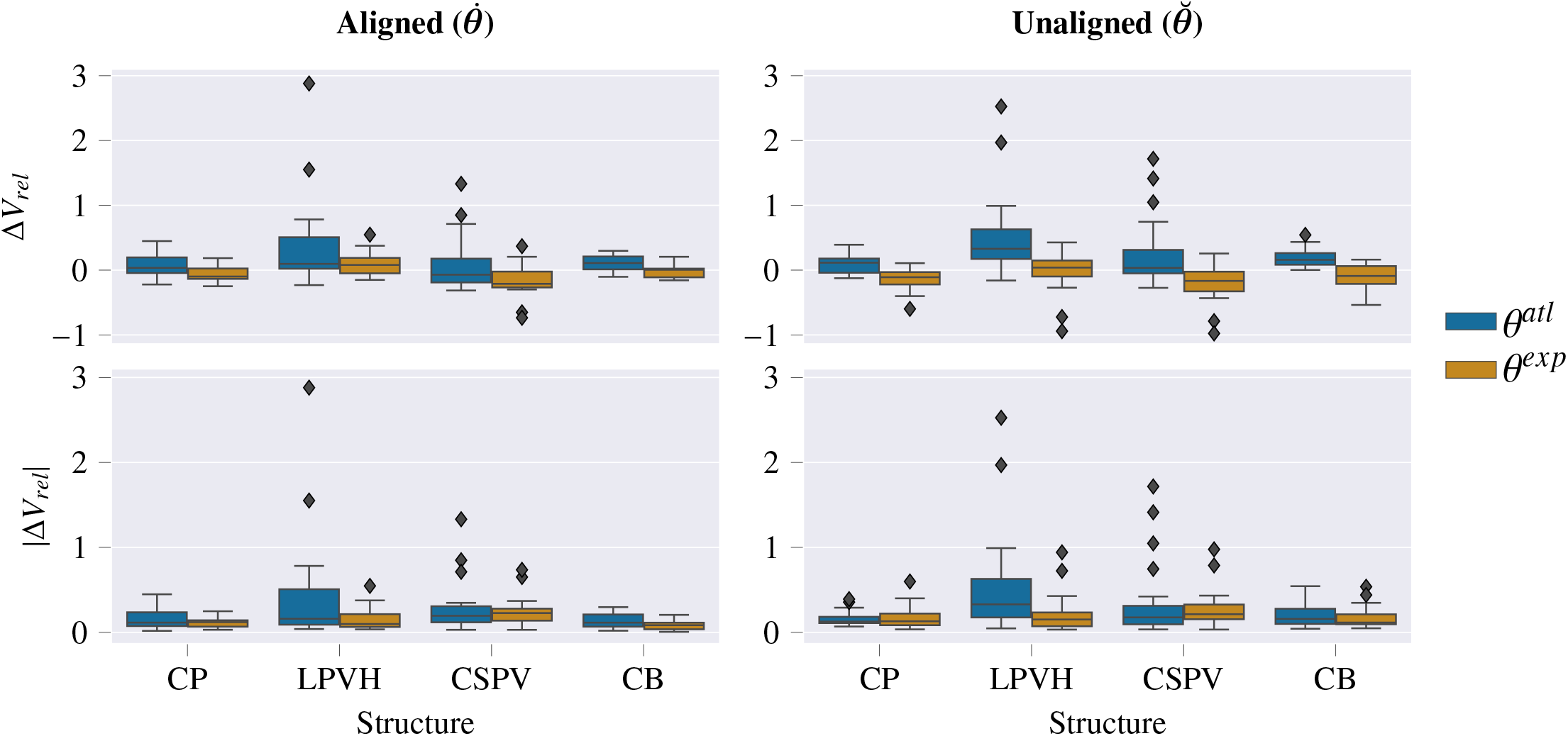
Signed and unsigned relative volume differences after post-processing for *θ^exp^* and *θ^atl^*.

#### Comparison to previous work

In Table 3, our resulting DSC performance using aligned images is compared to the intra-observer variability and to previous work performing subcortical structure segmentation. The intra-observariability was obtained by annotating ten images from the test set twice. Furthermore, for 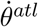 and *θ^exp^* the significance with respect to the naive propagated atlas labels is shown. The p-values shown were obtained by a repeated measures Analysis of Variance (ANOVA) for each structure, followed by post-hoc testing with a paired t-test. Reported p-values are from the post-hoc testing and underwent Bonferroni correction for the four structures, as well as for the three model comparisons (see Appendix C for full statistical results). For the CP, LPVH and CB, superior performance is obtained compared to previous work. The best performance obtained for these structures is also very close to the intra-observer variability in our test data (DSC of 0.85, 0.85 and 0.90 versus an intra-observariability of 0.86, 0.85 and 0.91 for the CP, LPVH and CB, respectively). For the CSPV the DSC is 0.03 lower than previously reported DSC scores, however, it should be noted that that study only performed segmentation in 2D standard planes [34].

#### Qualitative Results

In Fig. 6, an example prediction is shown for an aligned image at 22 GWs. The manual-ground truth as well as the predictions from both 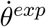 and 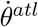 are shown. A qualitative inspection suggests that the predictions of the two networks closely resemble each other as well as the ground-truth. In Fig. 7, slices are shown containing the most prominent error modes in the aligned test set. To also visualize the segmented areas in the proximal hemisphere, the images in this figure are shown before any post-processing. From Fig. 7a-c, it can be observed that 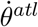 tends to oversegment in some areas, whereas 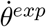 contains some spuriously segmented areas in the proximal hemisphere. Fig. 7d shows a CSPV segmentation at 18 GW with a very low DSC (0.51).

**Figure 6:**
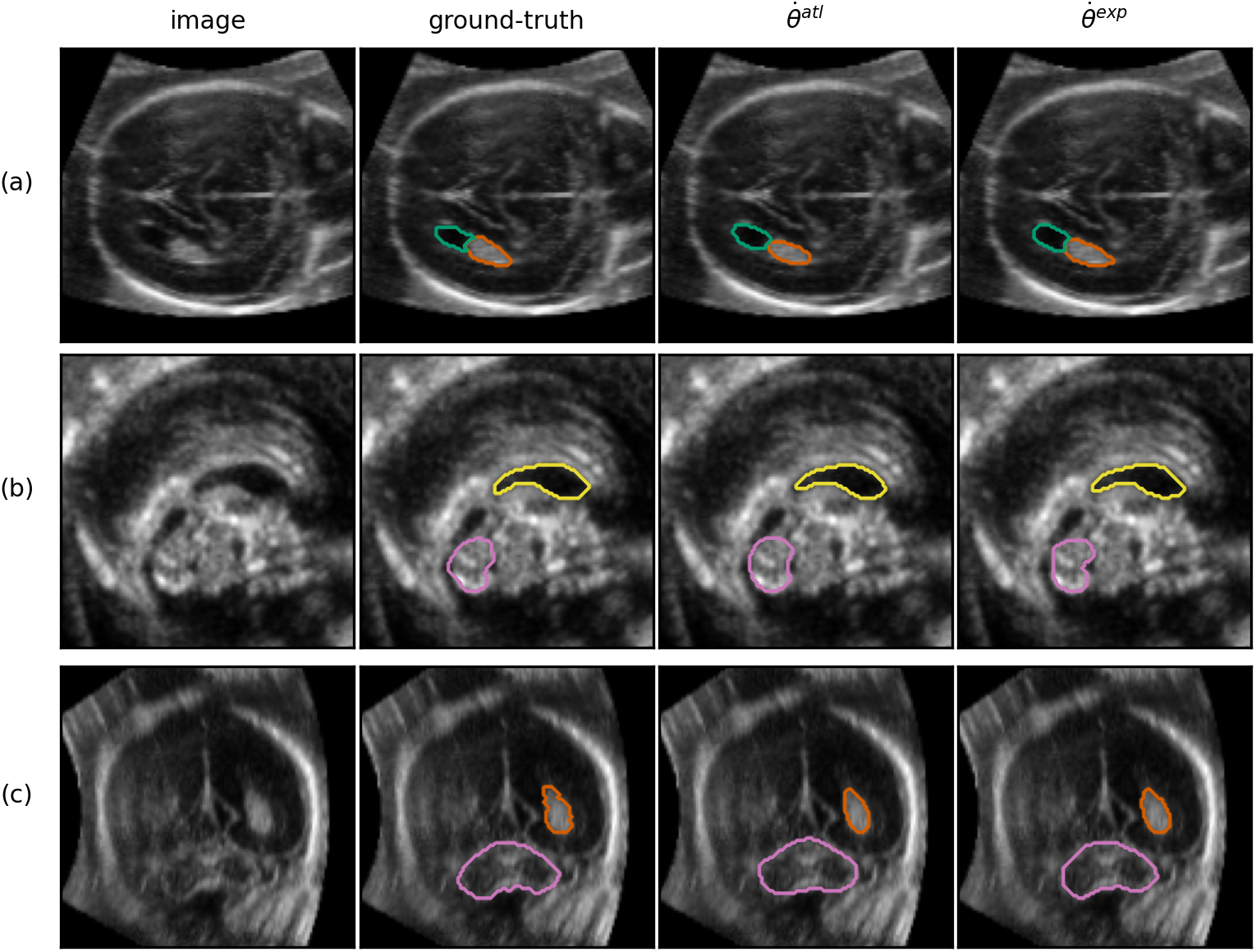
Example prediction of a test set volume at 22 GW (randomly chosen from 5 best performing images, average DSC across structures of 0.85), visualized in the axial (a), sagittal (b) and coronal (c) plane (segmentation was performed in 3D). The structures shown are the LPVH (green), CP (orange), CSPV (yellow), and the CB (pink). Only the outer boundary of the segmentation is shown in color for visualization purposes.

**Figure 7:**
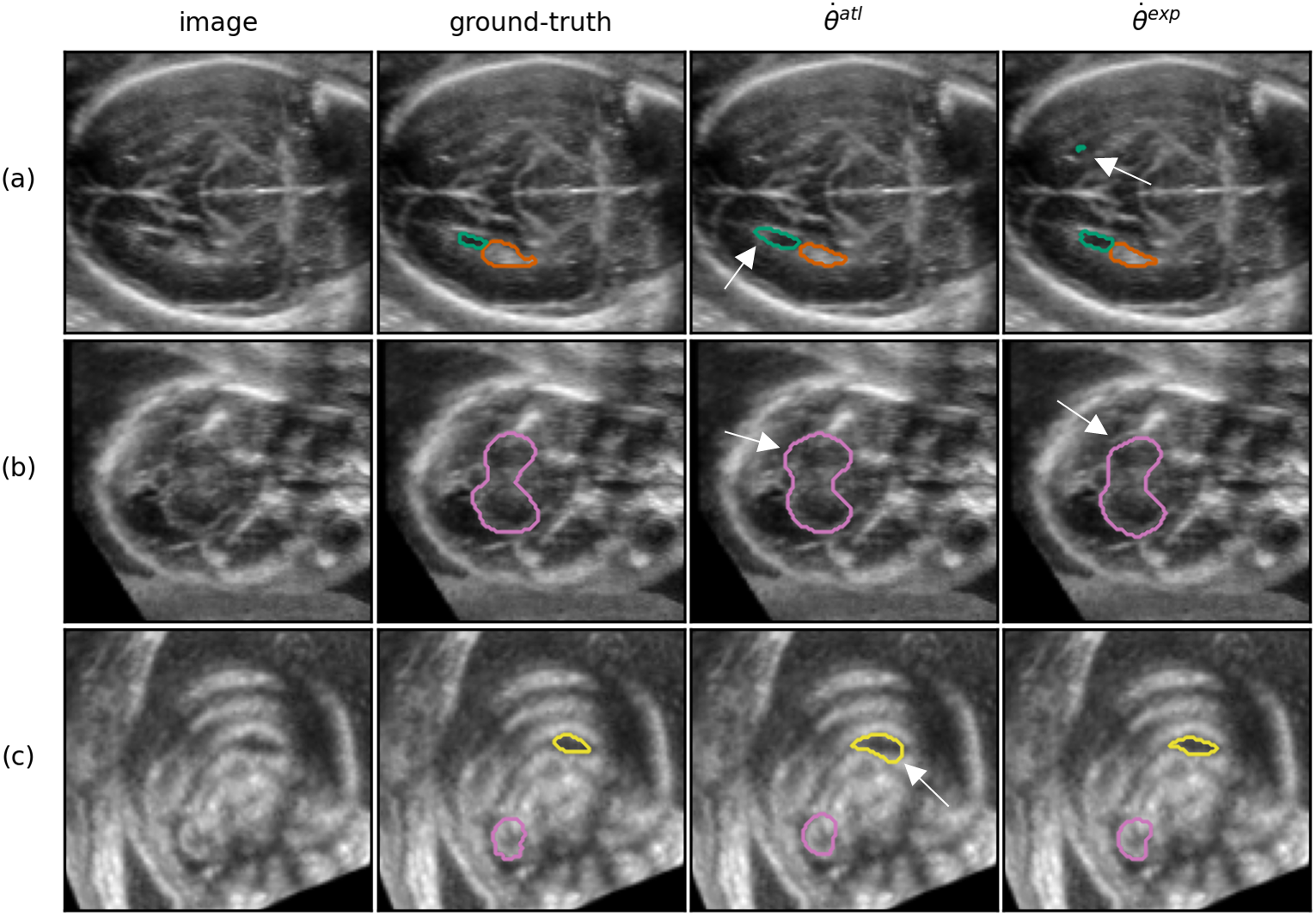
A set of examples showing the most prominent segmentation errors from 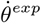 and 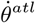 . In contrast to all other results in this section, the segmentations shown are before any post-processing so that false positive segmentation in the proximal hemisphere are visible. (a) axial plane of an image from a subject at 26 GW demonstrating oversegmentation of the LPVH (green) and LPVH segmentation in the distal hemisphere (b) axial plane of an image from a subject at 22 GW demonstrating over-segmentation of the CB (pink) for 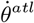 and slight under-segmentation for 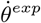 (d) sagittal plane of an image from a subject at 18 GW, demonstrating over-segmentation for 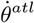 as well as the difficulty of manually segmenting the CSPV (yellow) at young ages.

### 4.3. Varying the number of manual annotations

For our second experiment, we reduced the amount of manual annotations, *n_a_*, required to generate the training data, thus effectively reducing the number of training images. The results of this experiment are shown in Fig. 8. It can be seen that for 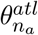, only a slight decrease in performance is observed when decreasing *n_a_*, both for the aligned (average DSC changed from 0.80 for *n_a_* = 9 to 0.78 for *n_a_* = 9) and unaligned images (average DSC changed from 0.78 to 0.76). For 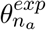, the performance decreases are stronger with lower numbers of training images, most notably in the unaligned scenario where the average DSC decreases to 0.33 for *n_a_* = 2.

**Figure 8:**
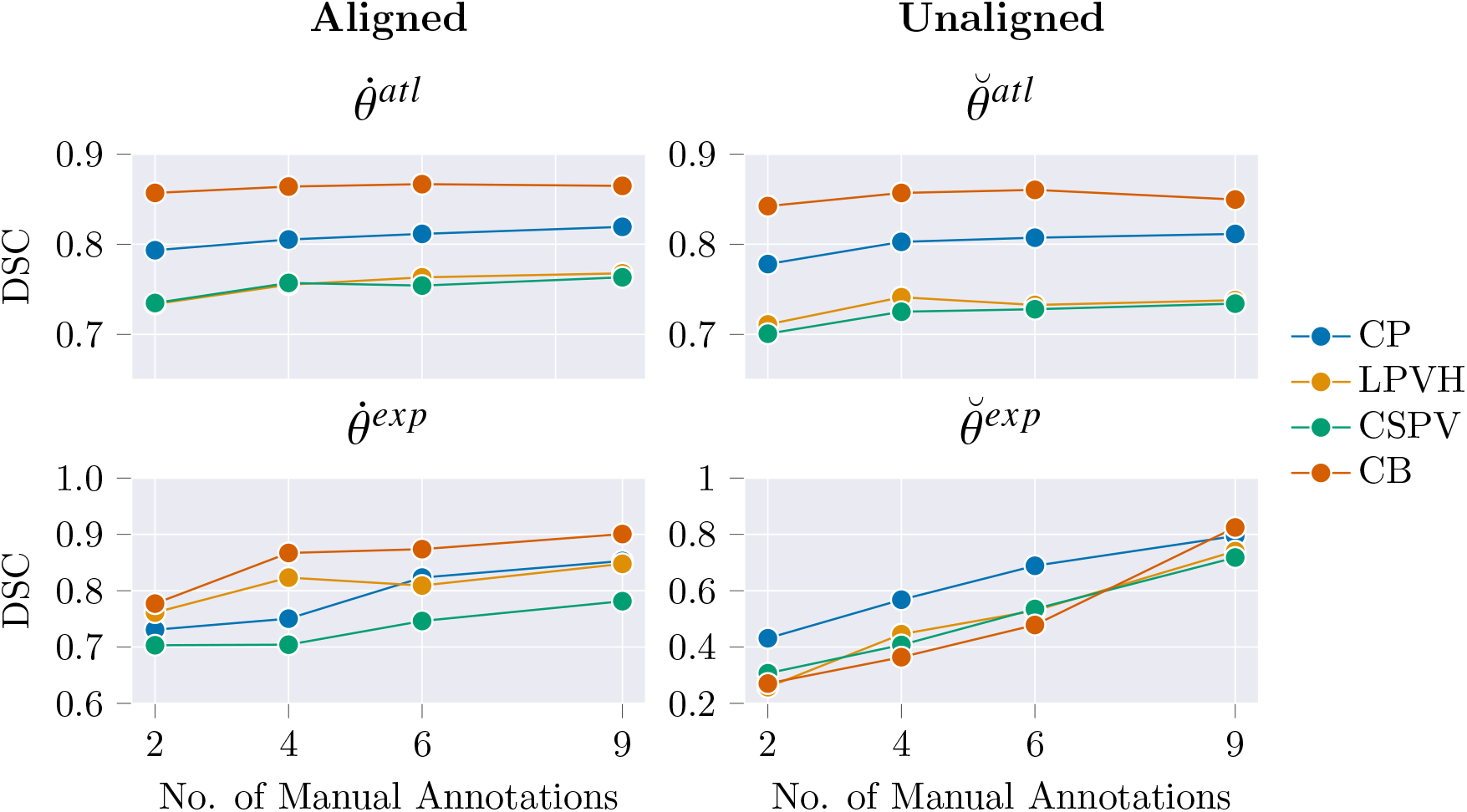
Varying the number of manual annotations (*n_a_*) used for generating the training labels.

### 4.4. Split training by GA

In Fig. 9, results are shown for networks trained separately for each twoweek gestational window, 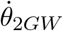, compared to having a single model for the whole age range, 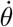, for both the atlas and expert labels. For 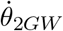, each point represents a single network trained using the images of the corresponding GW and the subsequent GW (i.e. 18 + 19), and is tested on the four images of the corresponding GW (i.e. 18), as test images are only available for every second week. For 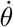, each point represents the performance on the four test images of the respective GW.

**Figure 9:**
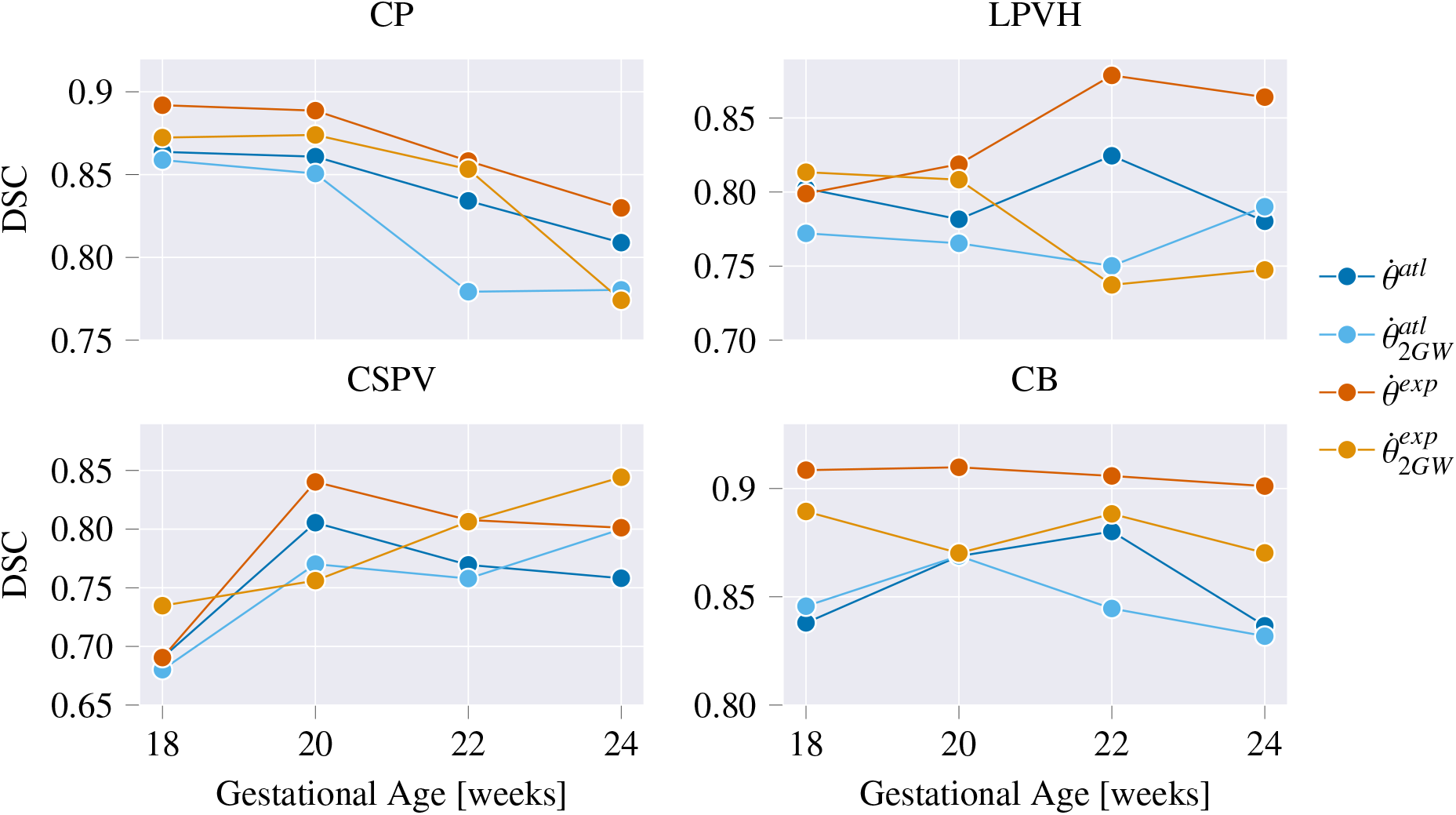
Performance of a single network 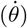, versus training separate models for every GA range of 2 weeks 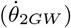.

It can be observed that the results for both label types, 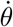 and 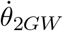, follow a similar trend. Averaged across all structures, 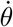 obtains a DSC of 0.81 and 0.85 for the atlas and expert labels, respectively, compared to a DSC of 0.80 and 0.82 for 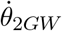 . For both the atlas and expert labels, 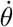 thus outperforms 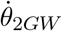 . This difference is significant (*p* < 0.05) for the CP, LPVH and CB in 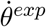, and only for the CP in 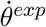 (p-values were obtained by a paired t-test with Bonferroni correction for the four structures).

From Fig. 9, the performance of 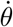 across gestation can also be observed. The CP shows a decrease of DSC with advancing GA whereas the CSPV segmentation performance increases. Both the CB and LPVH show a relatively stable performance across the GA range.

### 4.5. Clustered LPVH Labels

In Table 4 the LPVH segmentation performance of training 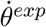 with atlas labels either propagated from 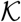 or 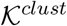 is shown. Training a network with the propagated clustered template annotations significantly improved the DSC performance from 0.69 to 0.77 (p<0.001) and *H*_95_ from 2.2 to 1.6 (p<0.05). Significance values were calculated using a paired t-test.

**Table 4:** LPVH segmentation performance using two different types of atlas-based annotation: LPVH labels obtained from annotating a single template per week 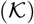 or annotating a set of clustered template images 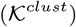.

| Network (template type) | DSC | $H_{95}$ [mm] |
| --- | --- | --- |
| $\dot{\theta}^{atl}(\mathcal{K})$ | 0.69 (0.13) | 2.2 (1.40) |
| $\dot{\theta}^{atl}(\mathcal{K}^{clust})$ | 0.77 (0.11) | 1.6 (1.2) |

### 4.6. Subcortical Growth Curves

The growth curves obtained from applying our trained networks are presented in Figs. 10 and 11. In Fig. 10 the predicted structural volumes are shown, whereas in Fig. 11 the predicted relative volumes with respect to the whole brain volume (*V_rel_brain_*), are shown. The growth curves were fitted with a linear or quadratic fit based on the significance of the quadratic component computed with a two-sided t-test. All parameters obtained through fitting are given in Appendix D.

**Figure 10:**
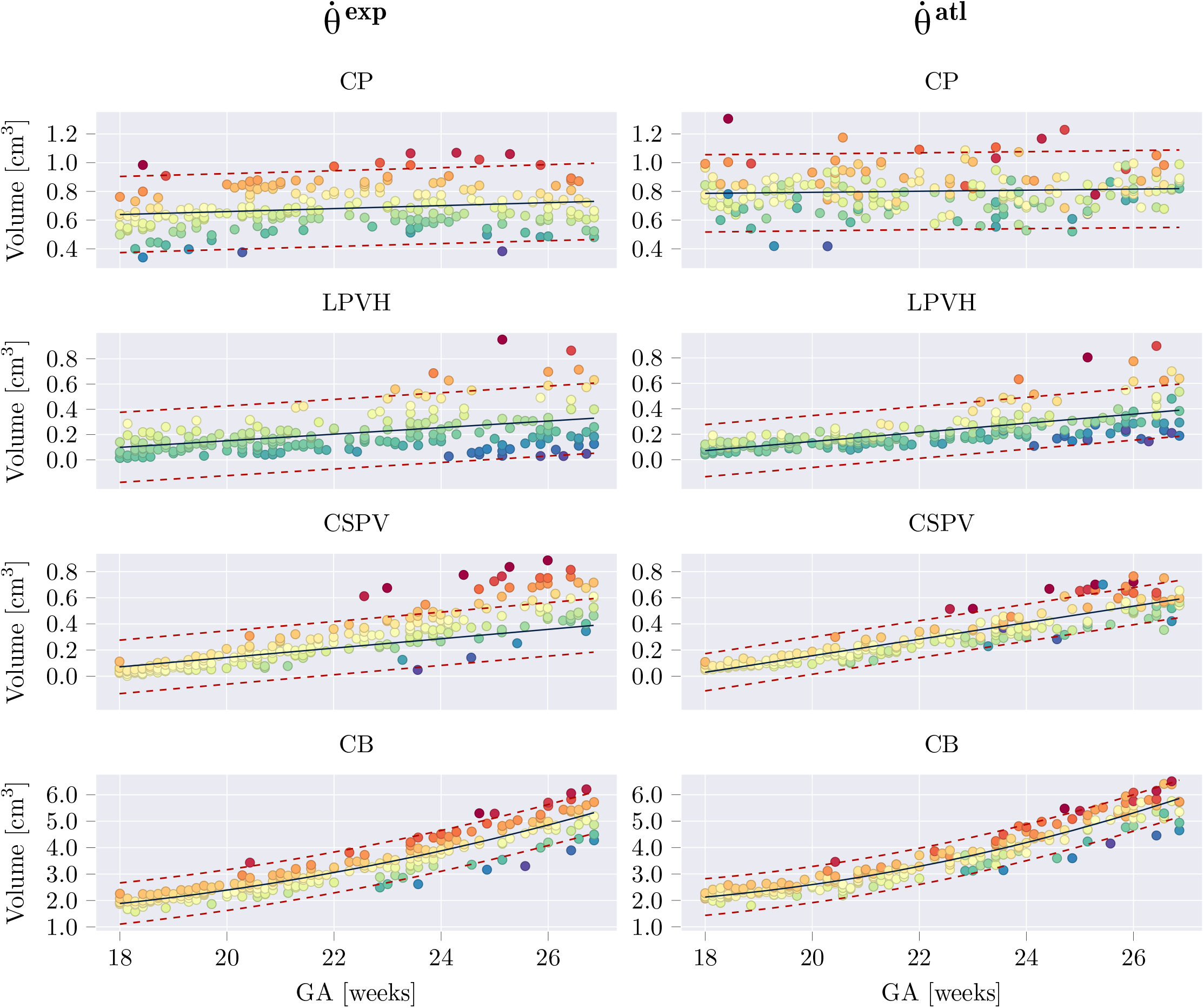
Estimated structural volumes for subcortical structures as a function of GA. Volumes were fitted with a linear or quadratic fit (black), in which the quadratic term was only added if it was significant for both networks (per structure). The 95% prediction confidence intervals where also computed and are shown with red dashed lines. For each structure, samples were colored based on their residual for 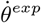, and the same colors (per sample) where used for 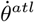.

**Figure 11:**
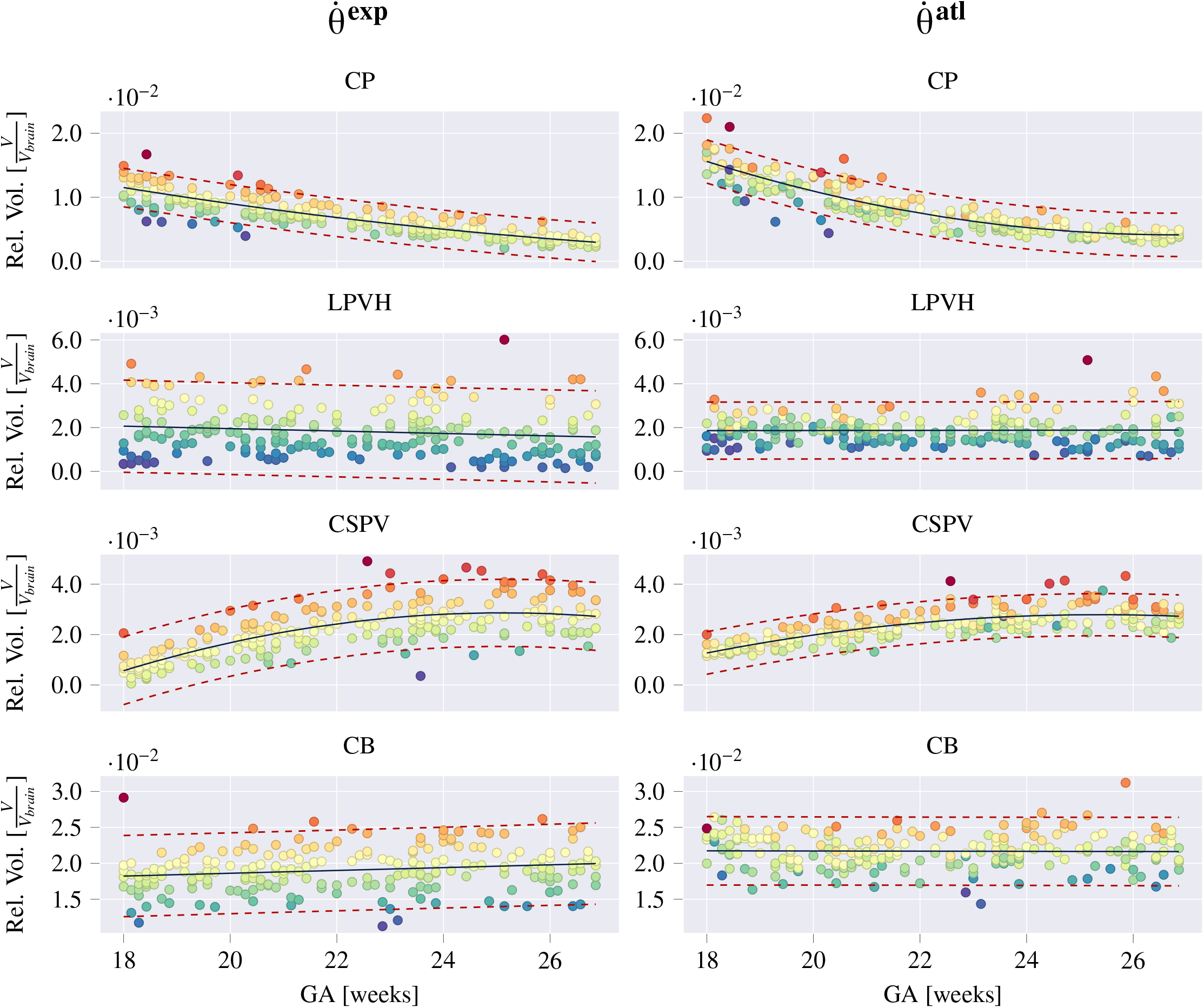
Estimated relative volumes (Rel. Vol.) with respect to the whole brain volumes (*V_rel_brain_*) for subcortical structures as a function of GA. Volumes were fitted with a linear or quadratic fit (black), in which the quadratic term was only added if it was significant for both models (per structure). The 95% prediction confidence intervals where also computed and are shown with red dashed lines. For each structure, samples were colored based on their residual for 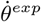, and the same colors (per sample) where used for 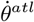.

To illustrate the level of consistency between 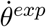 and 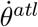, in Figs. 10 and 11 the samples were colored based on their residual value in the growth curves obtained from 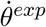. For a certain structure, samples with matching colors between the growth curves of the 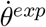 and 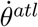 thus correspond to the same image. The choice of coloring the samples based on 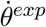 was made based on its better DSC performance (Table 3), but as it is only used to visualize the matching images, this choice is relatively arbitrary.

It can be observed that both of the models, 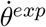 and 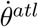, result in a similar growth trend and that the relative sample distribution is largely preserved (i.e. values above or below the growth curve are respectively above or below the curve for the other method, by a similar amount). However, 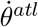 predicts larger volumes than 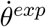 and there are also a few outliers that are not preserved between the two models.

As volumetric growth curves of the CB have been reported in previous work [16–22], we also compared our cerebellar growth curves to these studies (see Fig. B.15). This showed that our CB growth curves correspond well to previously reported values for cerebellar growth.

From the resulting growth curves (Figs. 10 and 11), it can be observed that the CB undergoes rapid growth during the second trimester of gestation, increasing from 1 cm^3^ at 18 GW to about 4 cm^3^ at the end of the second trimester. The CSPV shows a linear increase in structural volume during the studied GA range (from about 0.01 cm^3^ at 18GWs to 0.6 cm^3^ at 26 GWs) but its relative volume increases early in the second trimester and remains constant after approximately 24 GWs. The CP only shows a very small increase in structural volumes during the second trimester. However, the relative volumes show a rapid decline with respect to the total brain volume, ranging from 12% of the total brain volume at 18 GWs to 3% at 27 GWs. The LPVH structural volume increases linearly from about 0.1 cm^3^ to 0.3 cm^3^ in the studied GA range, whereas the relative volume shows a small downward trend.

## 5. Discussion

In this work, we performed subcortical segmentation in 3D fetal brain US images during the second trimester of gestation in a low-data regime. Our results showed that segmentation performance close to intra-observer variability can be obtained using only nine manually annotated brain images for training. We obtained best DSC performance of 0.85, 0.85, 0.78 and 0.90 for the CP, LPVH, CSPV and CB respectively, compared to an intra-observer variability of 0.85, 0.86, 0.86 and 0.91. We also applied our trained networks to generate longitudinal growth curves for subcortical structure development, that are, to the best of our knowledge, the first US-specific volumetric curves for the CP, LPVH and CSPV.

In our experiments, we investigated the impact of the number of labelled training samples, particularly assessing whether a network can be trained for structural segmentation in a low-data domain. We compared using a relatively large number (215) of images with weak labels (atlas labels) versus a very small number (9) of images with expert labels for training. For both settings, the number of images that had to be manually annotated to obtain the training labels was kept the same. Furthermore, we repeated our experiments for both *aligned* and *unaligned* images to explore the sensitivity of the model to brain pose. Somewhat surprisingly, in the aligned setting we obtained best performance using only 9 expertly annotated images, as opposed to a much larger number of weakly annotated images. This indicates that even though segmentation in 3D US by a human annotator is a very challenging task, a network is able to learn the correct features from only a small set of manual annotations, provided that the images are all aligned to the same coordinate space.

The network trained with expert labels showed a notably lower performance when trained with *unaligned* images than with the *aligned* images, which is most pronounced when decreasing the number of manual annotations used for training below nine (Fig. 8). On the other hand, this difference was not as pronounced for the network trained with atlas labels. This can be explained by the fact that in the *unaligned* setting the segmentation task consists of a combination of structure localization and subsequent segmentation. The weak, atlas labels contain imperfections at the boundaries of the segmentations, but are always approximately in the correct location. For this reason, these labels provide more information for the localization task than the few expertly annotated images and, as such, obtain better segmentation performance in the unaligned case. The few expertly annotated images in this setting are not sufficient to provide both localization and segmentation, which can most clearly be deduced from the large *H*_95_ distances (Fig. 4). These results thus suggest that a large amount of weak labels is better for localization, whereas few expert labels perform better if the task consists of solely delineating a structure in approximately the same location.

For the weak atlas labels, we observed only a small performance decrease when reducing the number of training images. Even by training with labels generated from just 2 GWs (coming from 2 manual annotations) the performance is close to the maximum performance (average DSC of 0.78 versus maximum DSC of 0.80 for the aligned case). The same pattern is observed for both *aligned* and *unaligned* data, confirming that the alignment does not considerably affect segmentation performance for the atlas labels. On the other hand, for the expert labels a performance decrease was observed when reducing the number of training images, showing a stronger decrease for the *unaligned* images than for the *aligned* images. As in this experiment we used as little as two images for training, while evaluating on images from the whole GA range, this drop in performance was in line with our expectations.

We also compared our resulting segmentation performance in the aligned setting to previous studies. However, it should be noted that performance metrics reported in different studies have to be carefully interpreted. Factors such as US image quality, structural definition (i.e. what is included in the segmentation), and GA range can all affect the difficulty of the task. The GA range affects the difficulty of the segmentation tasks as acoustic shadowing increases with advancing GA, due to calcification of the fetal skull. Furthermore, overlap measures, such as the DSC, generally report higher values for larger structures because a single (erroneous) voxel has a smaller effect on the resulting DSC value for these structures. We also want to stress that small performance increases are very difficult to quantify in 3D US segmentation due to the high intra- and inter-observer variability predominantly resulting from the ambiguity of the exact location of structural boundaries.

For the CP, LPVH, and CB we reported increased performance compared to previous studies (resulting in a DSC of 0.85, 0.85, and 0.90, respectively). An especially large increase was observed for the CB (DSC of 0.90 versus DSC of 0.80, 0.63 and 0.73 in previous studies). However, part of this increase can be attributed to the fact that in this study the CB was manually annotated including the bright echogenic boundaries, thus resulting in a larger segmented region, whereas this boundary was not included in [51] and [33]. This choice was made based on the fact that the transverse cerebellar diameter, which is clinically used to assess growth during a standard fetal examination, also includes this boundary [52]. Furthermore, manual segmentation is more consistent when including this boundary as the outside edge is generally clearly visible in the US images. For the CSPV, slightly higher performance (DSC of 0.81 versus 0.78) was reported in Huang et al. [34], but was obtained in the 2D midsagittal plane. Since the lateral boundaries of the CSPV are hardest to define in 3D images, this naturally results in larger prediction errors than 2D segmentations in this plane. Although these comparisons should thus be interpreted with care, it does show that we obtained competitive segmentation performance for all structures.

As the aim of developing a 3D segmentation model is to extract volumetric information, we also assessed the error of the resulting volume measurements in our test set (Fig. 5). These showed that the network trained with atlas labels generally produced positive volume differences, representing over-segmentation, indicating a tendency to segment a structure larger on the atlas than in the individual images. However, as the visualization of structural boundaries can be subjective in US, a consistent (albeit overestimating) segmentation prediction cannot be considered unacceptable.

We further analyzed our segmentation performance in the *aligned* setting by visualizing the most prominent errors before any post-processing (Fig. 7).

These results confirmed that the network trained with atlas labels showed a tendency to over-segment, whereas the network trained with expert labels resulted in some small segmented regions in the proximal hemisphere (as well as occasionally in other areas for the unaligned images). However, these small regions were easily removed by only retaining the largest connected component during post-processing. Furthermore, due to their small size, these regions

We thus believe that even though being trained with only a small number of manual annotations, both networks (trained with atlas or expert labels) perform exceptionally well on a task that is challenging for human annotators.

To observe the effect of training a single model for the entire GA range that was used in this study (18-26 weeks), we also trained separate networks for every biweekly GA window. These results showed that for most structures and GWs, the best performance is obtained by training a single model with data spanning the entire GA range. This confirms our expectation that information from all GWs provides longitudinal structural variability that improves the model’s representational power and, thus, segmentation performance. However, the differences in performance between both methods were small (DSC of 0.81 and 0.85 for the biweekly model versus a DSC of 0.80 and 0.82 for the single model for the atlas and expert labels, respectively), suggesting that models for a smaller GA window can also provide good performance.

In addition to comparing separately trained networks with a single model, Fig. 9 also shows the DSC performance as a function of the GW. The CSPV performance was low at 18 GWs and increased later in gestation, which matches our expectation that segmentation performance is better for larger structures, as the CSPV is very small (around 0.05 cm^2^ at 18 GWs versus 0.5 cm^2^ at 26 GWs) and therefore difficult to manually segment at 18 GW (Fig. 7d). The CP shows the opposite behavior, which can be attributed to the fact that the shape of the CP is more challenging to segment later in gestation but does not increase in size. Similarly, the CB is increasingly difficult to segment during gestation due to acoustic shadowing caused by increasing calcification of the temporal bone. However, as it also grows in size (from an average of 1 cm^3^ at 18 GW to 3.5 cm^3^ at 26 GW), the DSC performance is relatively stable across the GA range explored in this study. For the LPVH, both difficulty and size do not change drastically during gestation resulting in a consistent performance across the gestational period.

We extended our atlas-based labels with a clustering approach for the LPVH. This approach was only applied for the LPVH, as the other structures did not display as much shape variation within GWs. By annotating the clustered templates instead of a single template per GW, the DSC performance increased from 0.69 to 0.77 (*p* < 0.001), suggesting that the additional variation captured by the clustered templates contributed to an improved segmentation of the LPVH.

Finally, we applied our trained networks to extract volumetric information from a large number of images (n = 278), allowing us to generate US-specific growth curves for a healthy population. Previous work on volumetric subcortical development during gestation from US has largely been limited to cerebellar and thalamic growth [17, 39–41]. Therefore, to the best of our knowledge, this study provides novel US-specific volumetric growth curves of the CSPV, CP and LPVH for a geographically diverse, healthy population. We generated growth curves using the network trained with few expert labels as well as the network trained with atlas labels. This showed that between both networks the sample distribution was largely preserved, confirming that even though the network trained with atlas labels obtained lower performance due to over-segmentation, the predictions are largely consistent with the network trained with expert labels. Overall, this shows that both models can potentially be used to predict accurate subcortical volumes, however, care should be taken when comparing volumetric measures obtained from different models.

To validate the volumetric measures obtained in this study, we compared the growth curves of the structural CB volume to previously reported CB growth curves. This showed that our curves showed excellent agreement with past US studies, especially in the first half of the GA range. After 24 GWs the growth rates of the CB (slope of the growth curves) found in this study are slightly lower than in previous work. However, as described previously, the CB becomes more challenging to segment with advancing GA and, as such, this could explain the slightly deviating volume measurements at the end of the second trimester.

For the CB a rapid increase in volume was observed, whereas the relative volume, with respect to the total brain volume, remained mostly consistent (between 1.5% and 2.5 %). As the transverse cerebellar diameter is expected to increase linearly during the second trimester [1], this matches the quadratic growth of CB volume. The CP showed a rapid decrease in relative volume during the studied GA range. This aligns with the observation that early in the second trimester the CP almost completely fills the lateral ventricles and comprises a large part of the fetal brain, whereas later in gestation it appears as a small structure only partly filling the lateral ventricles. Structural volumes of the LPVH only showed a small increase during the second trimester, which is in agreement with the fact that the atrial width remains stable in healthy fetuses during the second trimester [1]. The structural CSPV volumes increase linearly, whereas the relative volumes show an increase in the first half of the studied GA range. As a deviating measurement of the CSPV has been related to agenesis of the corpus callosum [3] but is not widely studied, the presented growth curves might facilitate this in future work.

## 6. Conclusion

In summary, we showed that only a small number of annotated images are needed to successfully train a network for subcortical segmentation in 3D US images. To obtain optimal performance, alignment of the images is required beforehand, however, on unaligned testing images, a high performance can still be achieved using several weakly annotated images. By applying our trained networks to a large cohort of fetuses, we were able to generate US-specific growth trajectories of the CP, LPVH, CSPV and CB for a healthy population. This study thus demonstrates the feasibility of subcortical segmentation in 3D US using deep learning, and shows that volumetric measures obtained from these models can be used to obtain an improved understanding of subcortical growth during gestation. The models developed in this study offer the promise of comparing subcortical development between different fetal cohorts without doing time-consuming manual annotations in future studies. However, more work is needed to extensively validate the segmentation performance for datasets acquired with different US scanners as well as for cohorts containing pathological conditions.

## 7. Data and code availability statement

The image data is available from the INTERGROWTH-21^st^ Consortium upon reasonable request. All network training code as well as the weights of the trained networks can be requested by emailing the corresponding author. The groupwise registration algorithm used to generate the template image in this work was adopted from [46], and requests for this code can be addressed to the corresponding author of that paper.

## 8. CRediT authorship contribution statement

Linde Hesse: Conceptualization, Methodology, Investigation, Data Curation, Writing – original draft, Writing – review & editing, Software, Validation. Moska Aliasi: Data Curation, Writing – review & editing. Felipe Moser: Software, Writing – review & editing. Monique Haak: Writing – review & editing. Weidi Xie: Supervision, Writing – review & editing. Mark Jenkinson: Supervision, Writing – review & editing. Ana Namburete: Supervision, Conceptualization, Writing – Review & Editing, Funding Acquisition, Data Curation.

## 9. Acknowledgements

LH acknowledges the support of the UK Engineering and Physical Sciences Research Council (EPSRC) Doctoral Training Award. FM acknowledges the support and funding from the Engineering and Physical Sciences Research Council (EPSRC) and Medical Research Council (MRC) (EP/L016052/1), as well as the support from University College Oxford and its Oxford-Radcliffe benefaction. WX is supported by the UK Engineering and Physical Sciences Research Council (EPSRC) Programme Grant Seebibyte (EP/M013774/1) and Grant Visual AI (EP/T028572/1). MJ is supported by the National Institute for Health Research (NIHR) Oxford Biomedical Research Centre (BRC), and this research was funded by the Wellcome Trust [215573/Z/19/Z]. The Wellcome Centre for Integrative Neuroimaging is supported by core funding from the Wellcome Trust [203139/Z/16/Z]. AN is grateful for support from the UK Royal Academy of Engineering under the Engineering for Development Research Fellowships scheme, and to St Hilda’s College, Oxford.

We would also like to acknowledge the INTERGROWTH-21^st^ Consortium for collecting the data and making it available to us.

## Appendix A. List of Notations

**Table A.5:**
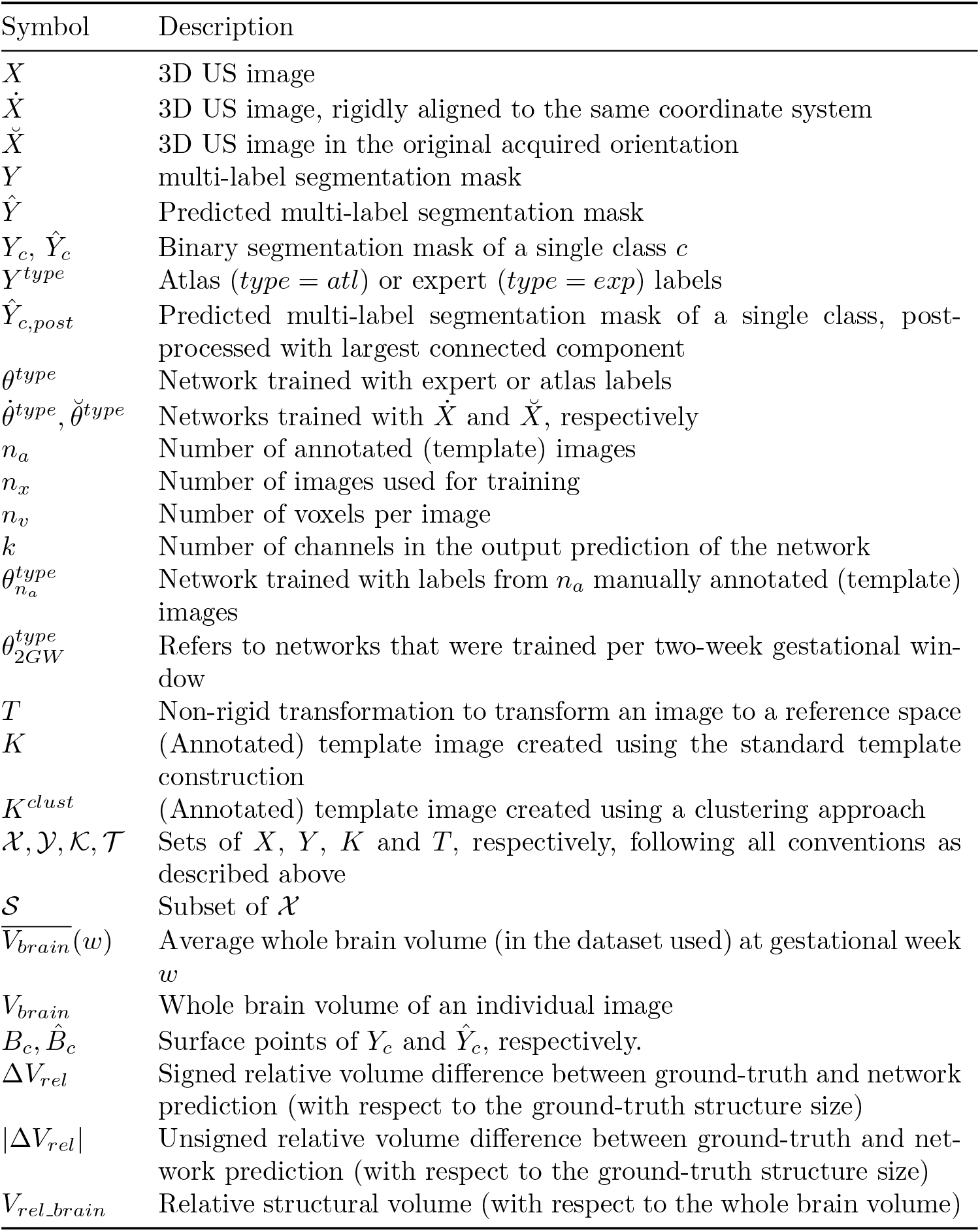
List of notations used in this work.

## Appendix B. Additional Figures

**Figure B.12:**
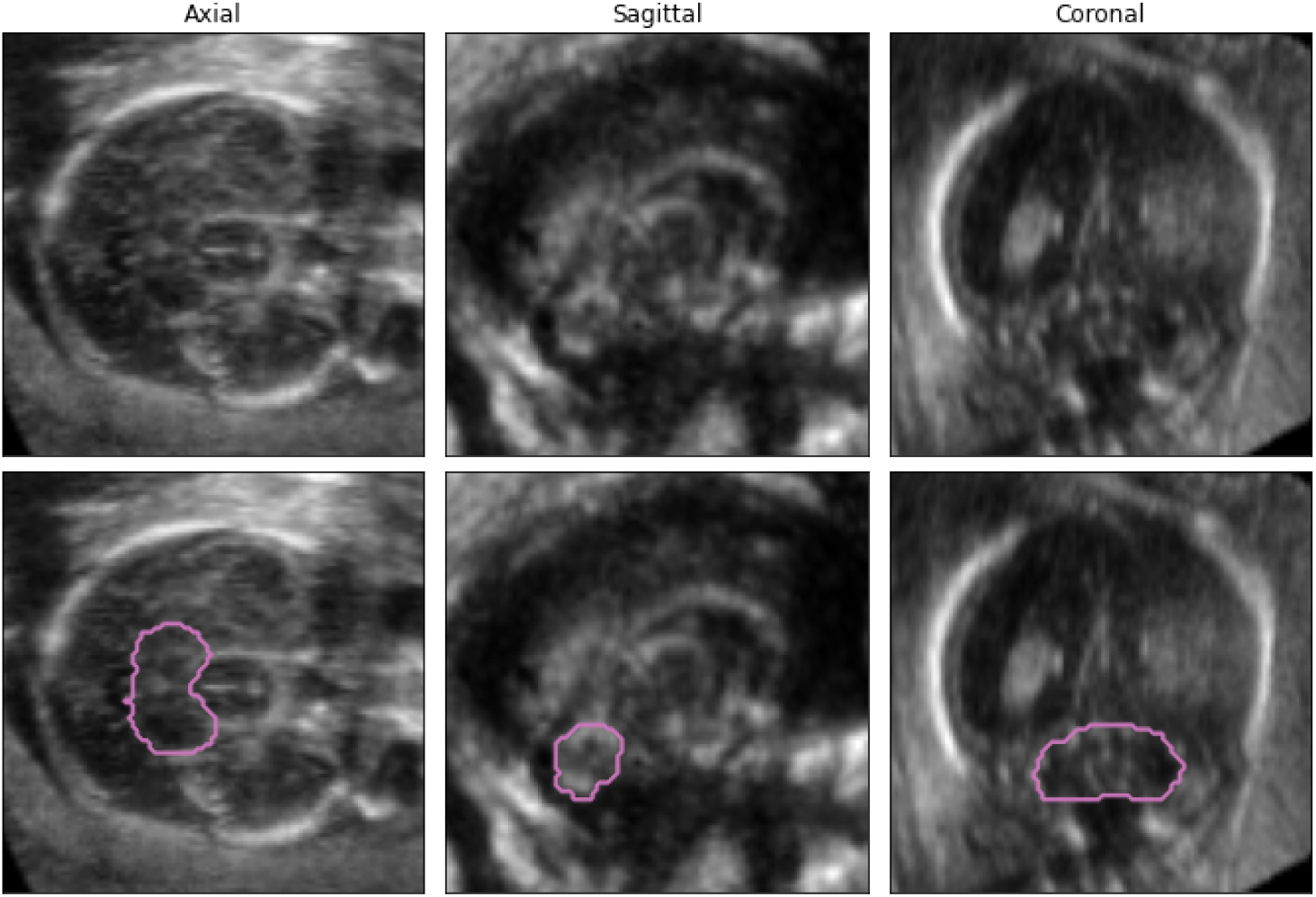
Example of a challenging manual annotation for the CB. The three columns each indicate one of the orthogonal 2D views. The top row shows the image without the annotation, and in the bottom row the outer boundaries of the manual labels are shown in pink on top of the image. It can be observed that for this image in both the axial and coronal plane the cerebellar boundaries are very hard to distinguish, whereas on the sagittal view the bean shaped CB is better visible.

**Figure B.13:**
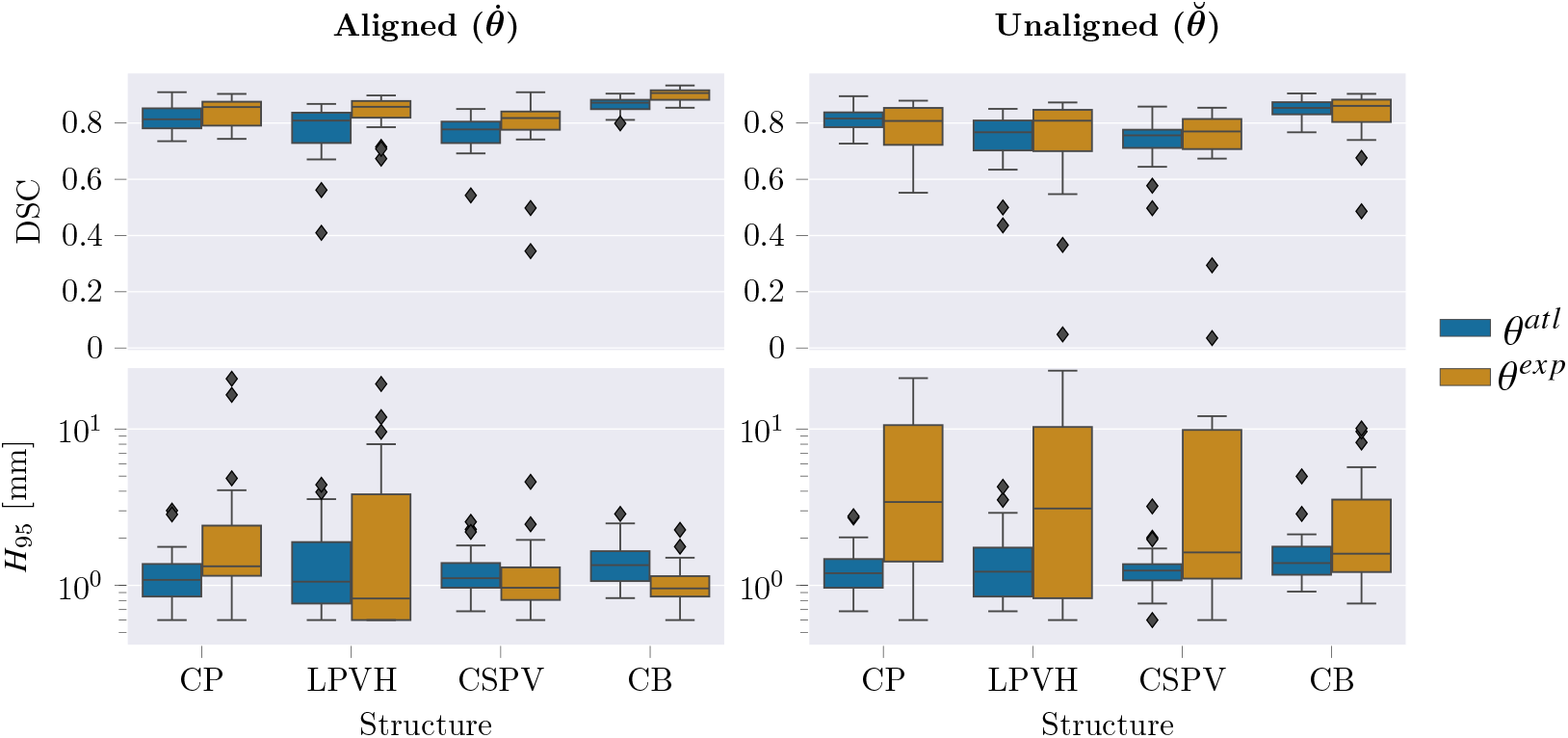
Resulting performance values before the largest connected component post-processing step was applied for *θ^atl^* and *θ^atl^*.

**Figure B.14:**
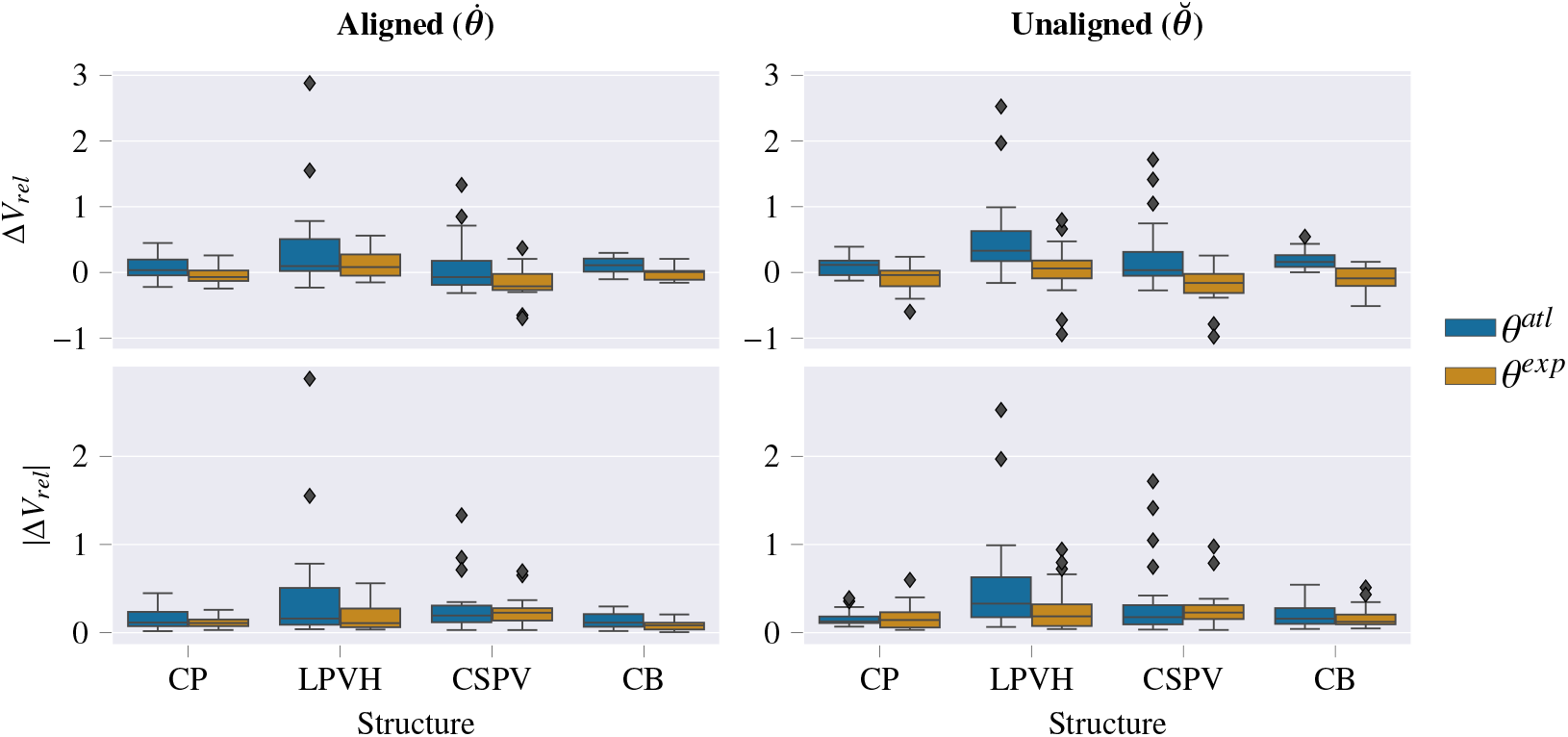
Volume-based performance values before the largest connected component post-processing step was applied for both *θ^atl^* and *θ^atl^*.

**Figure B.15:**
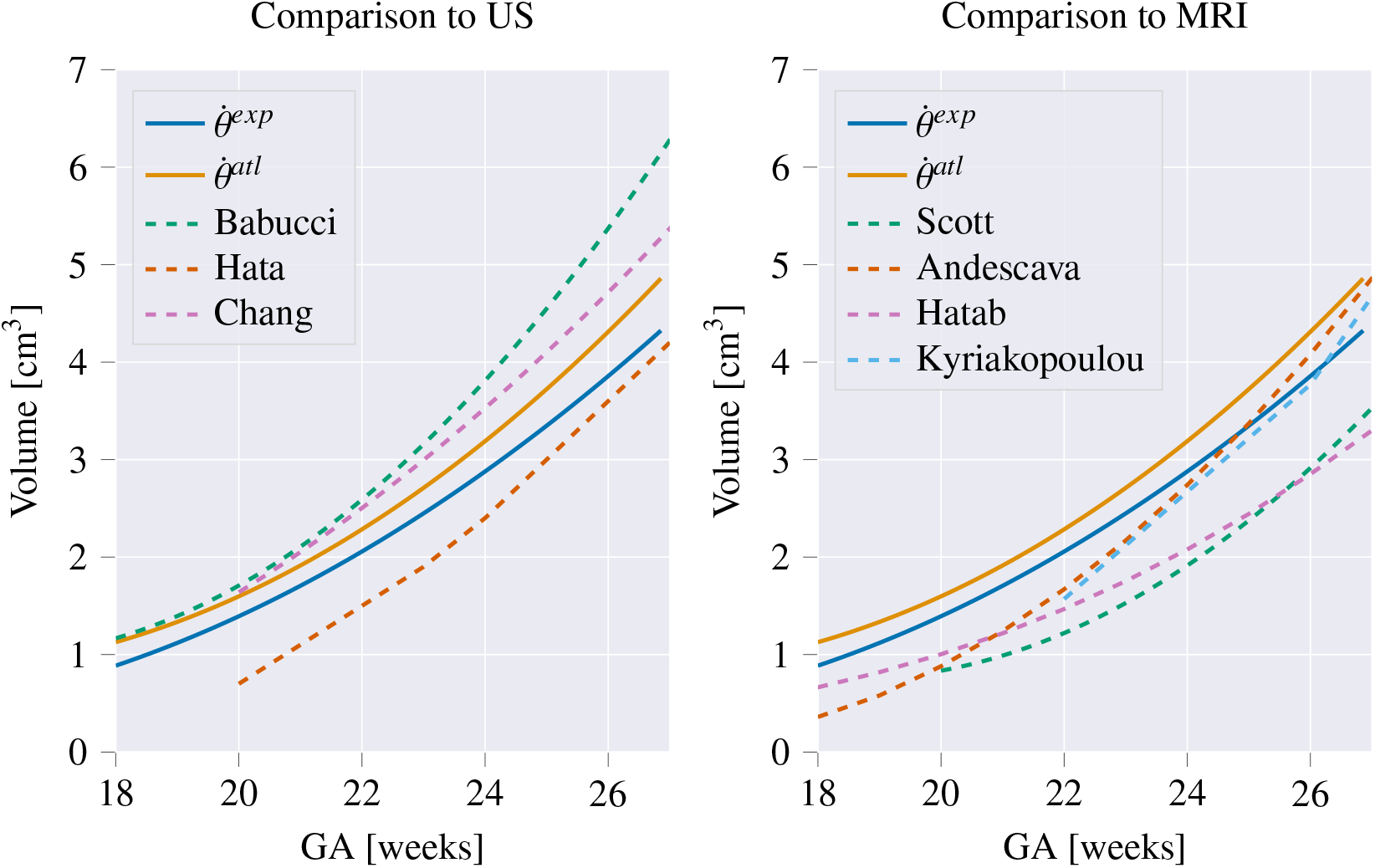
Comparison of growth curves obtained for the CB in this study to previous work. In the left panel the CB growth curves from our study (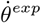 (blue) and 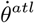 (orange)) are compared to other studies reporting cerebellar growth curves from US: Babucci [17] (green), Hata [21] (dark orange), Chang [18] (pink), and the right panel compares our curves to previously reported MRI growth curves: Scott [20] (green), Andescava [19] (dark orange), Hatab [22] (pink), and Kyriakopoulou [16] (light blue).

## Appendix C. Statistical Testing

**Table C.6:** P-values obtained from comparing the DSC segmentation performance of both networks, 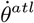 and 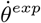, with each other as well as with the naive propagated atlas masks (*prop. atlas*). For each structure a repeated measures ANOVA was performed followed by post-hoc testing with a paired t-test. Reported p-values are post-hoc tests corrected with Bonferonni correction for the four structures, as well as for the three model comparisons. For the CSPV the ANOVA returned non-significant differences (p=0.08), and as such the post-hoc tests are not reported.

| | prop. atlas vs $\dot{\theta}^{atl}$ | atlas vs $\dot{\theta}^{exp}$ | $\dot{\theta}^{atl}$ vs $\dot{\theta}^{exp}$ |
| --- | --- | --- | --- |
| CP | <0.005 | <0.001 | <0.001 |
| LPVH | <0.001 | <0.001 | <0.01 |
| CSPV | - | - | - |
| CB | <0.05 | <0.005 | <0.001 |

## Appendix D. Growth Trajectories

**Table D.7:** Parameters resulting from the fitting of the structural volumes as function of the gestational age (*V* = *bias* + *w · x*1 + *w*^2^ *· x*2, with *w* the age given in weeks) for the different subcortical structures.

|  | model | bias | x1 | x2 | significance x2 |
| --- | --- | --- | --- | --- | --- |
| CP | expert | 0.7008 | 0.015 | -0.0054 | p < 0.001 |
|  | atlas | 0.8204 | 0.0125 | 0.0024 | p = 0.065 |
| LPVH | expert | 0.2072 | 0.0241 | -0.0004 | p = 0.79 |
|  | atlas | 0.2199 | 0.0354 | 0.0016 | p = 0.085 |
| CSPV | expert | 0.2857 | 0.0668 | 0.0009 | p = 0.38 |
|  | atlas | 0.2986 | 0.0610 | 0.0610 | p < 0.05 |
| CB | expert | 2.3112 | 0.3826 | 0.0197 | p < 0.001 |
|  | atlas | 2.6064 | 0.3924 | 0.0222 | p < 0.001 |

**Table D.8:** Parameters resulting from the fitting of the relative volumes (with respect to the whole brain volume) as function of the gestational age (*V_rel_brain_* = *bias*+*w·x*1+*w*^2^*·x*2, with *w* the age given in weeks) for the different subcortical structures.

|  | model | bias | x1 | x2 | significance x2 |
| --- | --- | --- | --- | --- | --- |
| CP | expert | 0.071 | -0.0010 | $4.654 \cdot 10^{-5}$ | $p < 0.005$ |
| | atlas | 0.084 | -0.0013 | 0.0001 | $p < 0.001$ |
| LPVH | expert | 0.0018 | $-7.785 \cdot 10^{-5}$ | $-5.397 \cdot 10^{-6}$ | $p = 0.6$ |
| | atlas | 0.0019 | $-4.979 \cdot 10^{-6}$ | $-4.017 \cdot 10^{-6}$ | $p = 0.5$ |
| CSPV | expert | 0.0021 | 0.0002 | $-4.697 \cdot 10^{-5}$ | $p < 0.001$ |
| | atlas | 0.0024 | 0.0001 | $-2.916 \cdot 10^{-5}$ | $p < 0.001$ |
| CB | expert | 0.0914 | 0.0001 | $-3.002 \cdot 10^{-5}$ | $p = 0.27$ |
| | atlas | 0.0224 | -0.0003 | $5.121 \cdot 10^{-5}$ | $p < 0.05$ |

